# Inter-organ communication shapes human metabolic tissue states and resolves anti-diabetic drug response modes in a six-tissue microphysiological system

**DOI:** 10.64898/2026.05.22.726943

**Authors:** Marissa McGilvrey, Shereen Chew, Mohd Farhan Siddiqui, Ronald Bronson, Merve Uslu, Shicheng Ye, Oscar Ospina, Andrea McPherson, Katrina Diel, Matjaz Dogsa, Luther Raechal, Martin Trapecar

## Abstract

Systemic glucose regulation depends on coordinated signaling among metabolically specialized tissues, yet most human *in vitro* models capture only limited portions of this network. Here, we developed and benchmarked a perfused human six-tissue MPS by combining AnthroHive, a recirculating perfusion platform, with MOTIVE-6, a six-compartment Multiorgan Tissue Interaction Vessel, to culture human gut epithelium, pancreatic islets, liver organoids, adipocytes, skeletal muscle, and midbrain-patterned brain organoids in a microphysiological system. Shared perfusion redirected engineered tissue states toward tissue-aligned metabolic, endocrine, absorptive, contractile, and neural-associated programs while reducing selected isolation-associated stress and remodeling signatures. Under High nutrient conditions, however, multi-tissue interaction shifted liver and islet responses toward inflammatory and nutrient-stress-associated gene expression, indicating context-dependent effects of cross-compartment signaling. Graded nutrient exposure resolved a staged circuit trajectory: Low nutrient conditions supported maintenance-associated programs, Mid nutrient exposure induced compensatory endocrine and anabolic remodeling with declining net glucose depletion, and High nutrient exposure shifted the system toward stress-associated metabolic dysfunction. Under High conditions, metformin and semaglutide produced distinct response modes. Metformin preserved circuit-level glucose handling without increasing insulin or C-peptide accumulation, while semaglutide remodeled gut, brain organoid, islet, and liver organoid transcriptional programs linked to nutrient sensing, epithelial maintenance, endocrine signaling, and neurometabolic state. Together, this study establishes a benchmarked human six-tissue MPS resource, paired with tissue-resolved transcriptomic, shared-media metabolomic, functional, endocrine, and inflammatory datasets, for investigating how tissue interaction, nutrient availability, and metabolic therapies reshape glucose-regulatory networks.

**Graphical Abstract:** 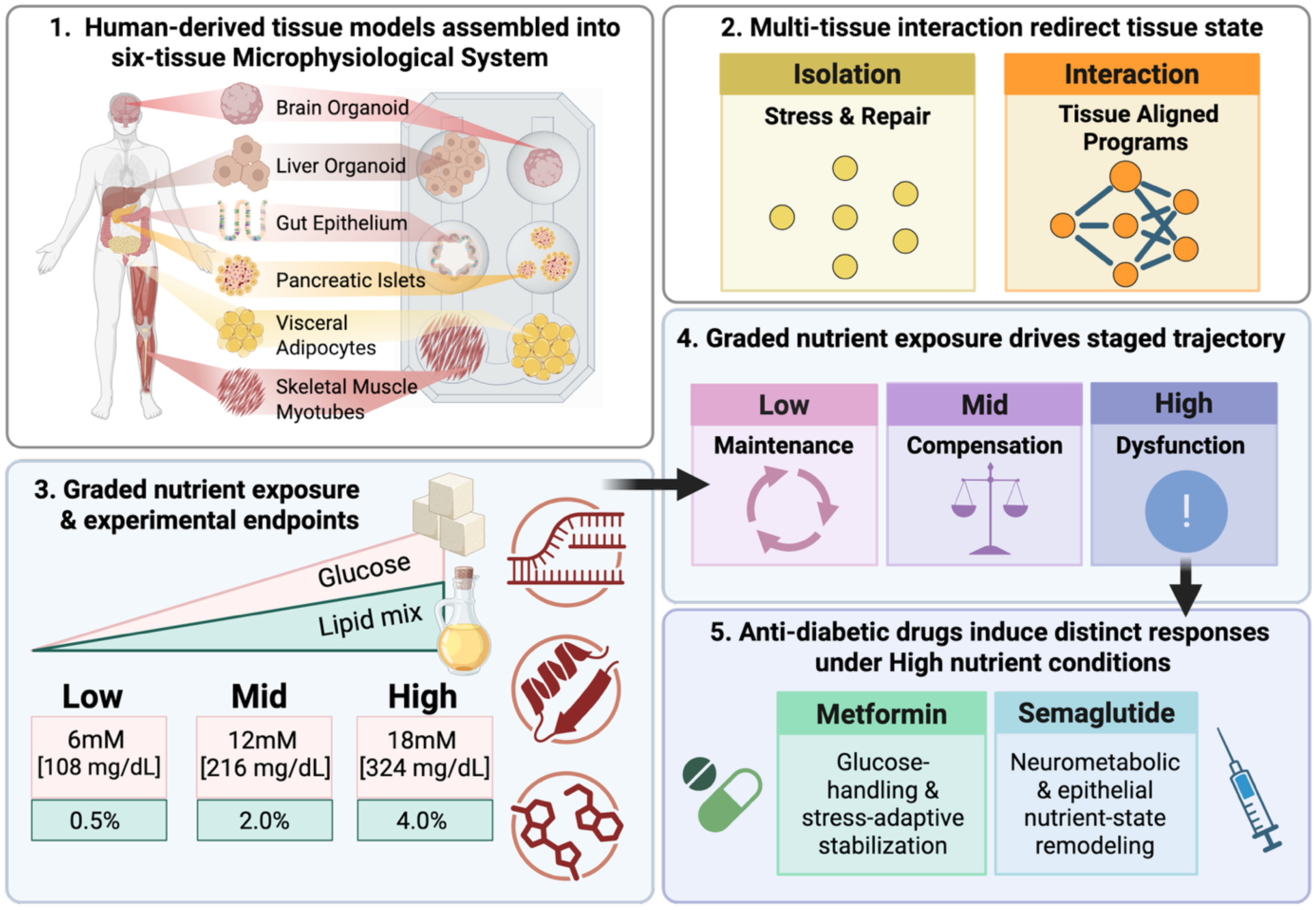

Created in BioRender. Trapecar, M. (2026) https://BioRender.com/a4tl7nv

## Introduction

Human tissues function within a circulating environment shaped by nutrients, hormones, cytokines, metabolites, trophic factors, and matrix-associated signals. These cues help regulate tissue state, tune functional output, and coordinate responses to changing physiological demand ^1^. Most human *in vitro* models remove tissues from this signaling environment. Organoids, primary cells, and engineered tissues may retain lineage-associated markers in isolation, but they lack the cross-compartment signaling that supports metabolic specialization and adaptive tissue function *in vivo.* This creates a need for human models that preserve tissue-specific readouts while reintroducing shared soluble signaling among metabolically relevant compartments.

This limitation is especially important for systemic glucose regulation. The gut, pancreatic islets, liver, adipose tissue, skeletal muscle, and brain each perform specialized metabolic functions, yet systemic metabolic state reflects their coordination ^1,2^. The gut epithelium regulates nutrient entry and enteroendocrine signaling, islets tune insulin and glucagon release, the liver balances glucose production with storage and lipid metabolism, adipose tissue and skeletal muscle contribute to fuel uptake and mobilization, and the brain responds to metabolic cues involved in energy-balance regulation ^3–8^. Responses to fasting-like states, increased post-prandial nutrient availability, and sustained nutrient overload therefore involve coordinated multi-tissue remodeling beyond changes in substrate availability. A model intended to study distributed human glucose-regulatory tissue states may therefore benefit from incorporating absorptive, endocrine, hepatic, storage, oxidative, and nutrient-sensing compartments within the same circulating environment.

A central challenge in human metabolic biology is understanding how distributed tissue networks coordinate adaptive responses to changing nutrient availability, and how these same networks become dysregulated during chronic metabolic stress ^1,2^. In response to changing nutrient availability, tissues can adapt through altered substrate handling, endocrine signaling, mitochondrial remodeling, and transcriptional regulation. With sustained nutrient overload, these responses can shift toward insulin resistance, inflammatory remodeling, impaired glucose handling, and energetic dysfunction ^2,9–11^.

This transition contributes to type 2 diabetes, metabolic syndrome, and age-associated metabolic decline, and has been linked to neurodegeneration-relevant changes in brain energetics and systemic metabolic signaling ^12–17^. The human tissue interactions that support functional state, and the contexts in which shared signaling contributes to stress propagation, remain difficult to study experimentally ^1,7,18^.

Existing models have advanced understanding of systemic metabolic regulation, but it remains difficult to experimentally isolate how human tissue interactions shape metabolic state across defined nutrient and therapeutic contexts. Isolated human cultures enable mechanistic control but lack circulating signals that shape metabolic state. Animal models preserve organism-level physiology but introduce species-specific differences in endocrine regulation, lipid handling, glucose set points, neuro-metabolic coupling, and therapeutic response ^19–21^. Microphysiological systems (MPS) offer a complementary strategy by reconstructing selected features of human physiology under controlled conditions ^21,22^. Co-culture and organ-on-chip models have advanced the study of metabolic crosstalk, including liver–pancreas, gut–liver, adipose–liver, and liver–islet interactions ^21,23–32^. Recent human MPS studies further show that perfused tissue-axis models can resolve selected features of systemic-milieu-dependent remodeling, including adipose–liver crosstalk, inflammatory and metabolic dysfunction, and therapeutic modulation within short experimental windows ^33^. However, systemic glucose regulation spans a broader network of nutrient-sensing, endocrine, storage, oxidative, inflammatory, and neuro-metabolic tissues. What remains less established is whether a human multi-tissue MPS can resolve coordinated responses to defined nutrient and therapeutic perturbations using tissue-resolved molecular and shared-media functional readouts. This is particularly relevant for metabolic therapies, which can act through hepatic, intestinal, endocrine, and central mechanisms and may produce responses that are difficult to assign from single-tissue readouts alone ^34,35^.

Here, we developed and benchmarked a compartmentalized shared-circulation human six-tissue MPS as a resource for studying glucose-regulatory tissue interaction. Using AnthroHive and MOTIVE-6, we assembled intestinal epithelium, pancreatic islets, liver organoids, visceral adipocytes, skeletal muscle, and midbrain-patterned brain organoids within a shared circulating media environment while preserving tissue-specific recovery for downstream analysis. We paired functional assays, endocrine and inflammatory profiling, bulk RNA-seq, and untargeted shared-media metabolomics to define how tissue interaction, graded nutrient availability, sustained nutrient overload, and anti-diabetic drug perturbation reshape tissue and circuit-level states. This study establishes a reusable experimental and analytical framework for interrogating cross-compartment signaling in human metabolic regulation and provides a public multi-omics dataset for future analysis.

## Results

### AnthroHive and MOTIVE-6 establish a configurable shared-perfusion platform for six-tissue metabolic MPS studies

To model selected features of distributed human glucose regulation, we mapped six metabolically relevant tissue modules onto a shared-circulation MPS architecture: intestinal epithelium, liver organoids, pancreatic islets, visceral adipocytes, skeletal muscle, and midbrain-patterned brain organoids (Fig.1A,B). To implement this architecture, we developed AnthroHive, a custom multi-vessel perfusion platform designed to support interconnected microphysiological system modules under continuous recirculating flow (Fig.1C). AnthroHive can house up to nine MPS vessels simultaneously and provides tunable perfusion across a controllable range of 0.5–3 mL/min, regulated through an integrated e-paper display. The platform supports customizable vessel configurations, including the Multiorgan Tissue Interaction Vessel-6 (MOTIVE-6; Fig.1B,D), a six-compartment vessel designed to culture distinct tissue types within a shared circulating media environment. MOTIVE-6 accommodates six individual 24-well-format membrane inserts, allowing selected tissue compartments to be incorporated or exchanged depending on the biological question. Each tissue is cultured on a 0.4 µm pore-size permeable membrane insert, allowing soluble metabolites, hormones, cytokines, and other secreted factors to enter the shared perfusate while maintaining physical separation between tissue compartments. This design enables controlled shared-media exposure and soluble cross-compartment signaling without direct tissue contact while preserving compartment-specific tissue recovery for downstream molecular and functional profiling. For all six-tissue experiments, MOTIVE-6 compartment positions were held constant across vessels and biological replicates, preserving a defined flow path and spatial arrangement for tissue-specific analysis within the interacting circuit (Fig.1B). Thus, AnthroHive and MOTIVE-6 provide a compartmentalized shared-circulation platform for testing whether inter-tissue communication reshapes engineered human metabolic tissue state.

**Figure 1.**
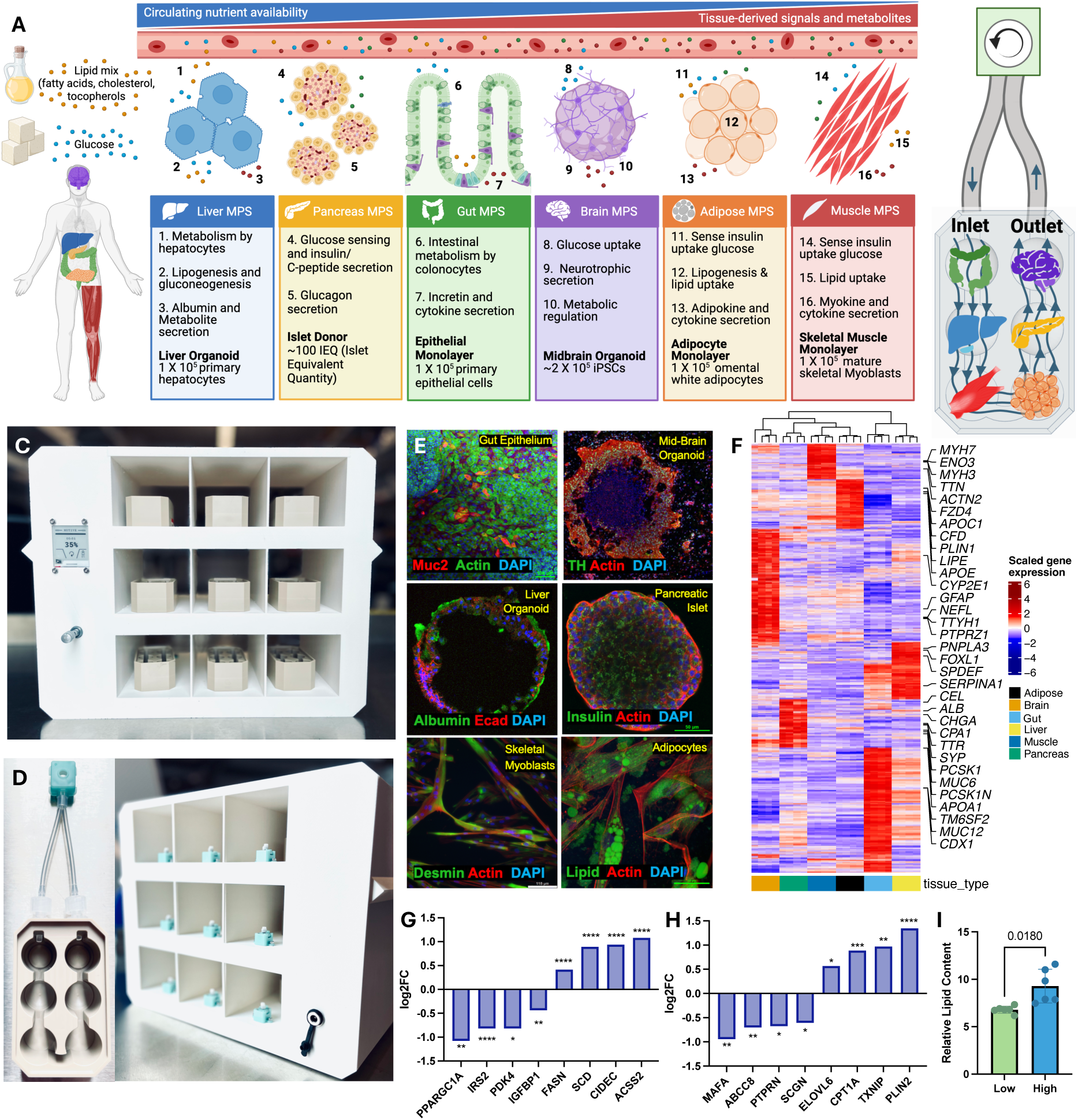
AnthroHive and MOTIVE-6 establish a perfused human six-tissue metabolic circuit. **(A)** Overview of the six human tissue compartments selected to model major features of systemic glucose and lipid regulation: gut epithelium, liver organoids, pancreatic islets, visceral adipocytes, skeletal muscle, and midbrain-patterned brain organoids. **(B)** Spatial arrangement of tissue compartments within MOTIVE-6: gut, green; liver organoids, blue; skeletal muscle, red; visceral adipose tissue, orange; pancreatic islets, yellow; brain organoids, purple. **(C)** AnthroHive multi-vessel perfusion platform used for controlled flow and parallel culture of six-tissue MPS. **(D)** MOTIVE-6 culture vessel with perfusion pump head and tubing used to maintain six tissue modules under shared-media circulation. **(E)** Immunofluorescence characterization of tissue phenotypes: gut epithelium, MUC2; midbrain-patterned brain organoids, tyrosine hydroxylase (TH); liver organoids, albumin; pancreatic islets, insulin; skeletal muscle myotubes, desmin; visceral adipocytes, intracellular lipid accumulation. **(F)** Bulk RNA-seq of tissue modules maintained in isolation. Heatmap shows selected tissue-enriched genes identified by differential expression analysis across isolated tissue modules; log(FC) >1, FDR <0.05 with Benjamini-Hochberg correction. **(G)** Liver organoid and **(H)** pancreatic islet transcriptional responses to Low versus High nutrient exposure in isolation. Low and High indicate 6 mM glucose/0.5% Lipid Mixture 1 and 18 mM glucose/4% Lipid Mixture 1, respectively. Bars show DESeq2-derived log2 fold change for selected nutrient-responsive genes; positive values indicate higher expression under Low conditions and negative values indicate higher expression under High conditions. n = 3 for isolation and n = 6 for interaction for each tissue module. Significance indicates Benjamini-Hochberg-adjusted P values from the RNA-seq differential expression model; *P < 0.05, **P < 0.01, ***P < 0.001, and ****P < 0.0001. **(I)** Quantification of intracellular lipid accumulation in mature visceral adipocytes cultured under Low versus High nutrient conditions, calculated as LipidTOX intensity normalized to nuclei count. n = 6 biological replicates; each replicate represents the average of six images. Statistical significance was determined by unpaired t-test.

### Engineered tissue modules retain tissue-associated identity and nutrient responsiveness

Before testing multi-tissue interaction, we benchmarked each tissue module in isolation to confirm tissue-associated identity, functional features, and baseline nutrient responsiveness under the culture conditions used for subsequent shared-perfusion experiments. Large intestinal epithelium, liver organoids, donor-derived pancreatic islets, visceral adipocytes, skeletal muscle cultures, and midbrain-patterned brain organoids were evaluated using immunofluorescence, bulk RNA-seq, and selected functional or phenotypic assays, including TEER, albumin secretion, and lipid accumulation where applicable (Fig.1E,F; Fig.S1).

Bulk RNA-seq of isolated tissue modules showed distinct tissue-associated gene expression profiles, with each compartment clustering by enriched transcript patterns in the heatmap before shared-perfusion studies (Fig. 1F). Large intestinal epithelial cultures expressed MUC2 and intestinal-associated transcripts including *MUC12*, *CDX1*, and *TM6SF2*, supporting epithelial identity and barrier-associated programs (Fig.1E,F). Consistent with barrier-forming capacity, intestinal epithelial cultures developed high TEER in isolation before integration into shared perfusion experiments (Fig.S1A). Midbrain-patterned brain organoids expressed tyrosine hydroxylase and neural/glial-associated transcripts including *GFAP*, *NEFL*, *TTYH1*, and *PTPRZ1,* supporting brain organoid-associated transcriptional features (Fig.1E,F). Visceral adipocytes displayed lipid droplet accumulation and expressed adipocyte-associated lipid-handling genes including *PLIN1*, *APOE*, *APOC1*, *CFD*, *FZD4*, and *LIPE* (Fig.1E,F). Skeletal muscle cultures expressed desmin and structural muscle transcripts including *ACTN2*, *TTN*, *MYH7*, and *MYH3*, supporting myogenic and contractile-associated programs (Fig.1E,F). Liver organoids retained hepatic/hepatobiliary-associated features, with expression of albumin and transcripts including *ALB*, *CYP2E1*, *SERPINA1*, *PNPLA3*, *FOXL1*, and *SPDEF*, as well as detectable albumin secretion after differentiation (Fig.1E,F; Fig.S1C). Donor-derived pancreatic islets exhibited insulin staining and expressed endocrine and pancreatic-associated genes including *PCSK1N*, *PCSK1*, *CHGA*, *CEL*, *TTR*, and *CPA1* (Fig.1E,F). These protein marker, transcriptional, and functional readouts confirmed tissue-associated features across the six modules before shared-perfusion studies.

We next assessed whether selected tissue modules retained baseline responsiveness to the Low and High nutrient conditions used in the experimental series. In isolated liver organoids, High versus Low nutrient exposure increased expression of genes linked to mitochondrial regulation, insulin-responsive signaling, nutrient oxidation, and stress-responsive growth factor signaling, including *PPARGC1A*, *IRS2*, *PDK4*, and *IGFBP1* (Fig.1G). Low exposure increased genes associated with lipogenesis, fatty-acid desaturation, lipid-droplet storage, and acetate-dependent lipid metabolism, including *FASN*, *SCD*, *CIDEC*, and *ACSS2* (Fig.1G), indicating nutrient responsiveness without a simple fasting-like oxidative signature. Islets showed a similar nutrient-responsive shift. High exposure increased β-cell identity and secretory-function genes, including *MAFA*, *ABCC8*, *PTPRN*, and *SCGN*, whereas Low exposure increased lipid-metabolism, fatty-acid-handling, and stress-responsive nutrient-sensing genes, including *ELOVL6*, *CPT1A*, *TXNIP*, and *PLIN2* (Fig.1H). Visceral adipocytes accumulated more intracellular lipid under High compared with Low nutrient exposure, confirming phenotypic responsiveness to nutrient excess (Fig.1I; Fig.S1D).

Together, these validation studies established baseline tissue identity and nutrient responsiveness for interpreting subsequent circuit-level responses to shared perfusion and graded nutrient exposure.

### Multi-tissue interaction shifts engineered tissues from isolation-associated maintenance states toward tissue-specific functional programs

With previous human physiomimetic studies, we have demonstrated that connecting tissue modules can reshape inflammatory and metabolic programs across linked compartments. Gut–liver physiomimetics revealed context-dependent modulation of IBD-associated inflammation by microbiome-derived short-chain fatty acids ^36^, while a gut–liver–brain model showed that circulating metabolites and inflammatory signals can propagate across connected human tissue systems relevant to neurodegenerative disease biology ^37^. These studies established tissue interaction as an active experimental variable that can alter engineered tissue state. Building on this framework, we next tested whether assembling six validated human tissue compartments into shared perfusion altered engineered tissue state across a broader metabolic circuit (Fig. 2A). Isolated and interacting cultures were compared after 4 days under the Mid nutrient condition to separate the effect of tissue interaction from nutrient-dose effects and to establish the reference state for subsequent perturbation studies.

**Figure 2.**
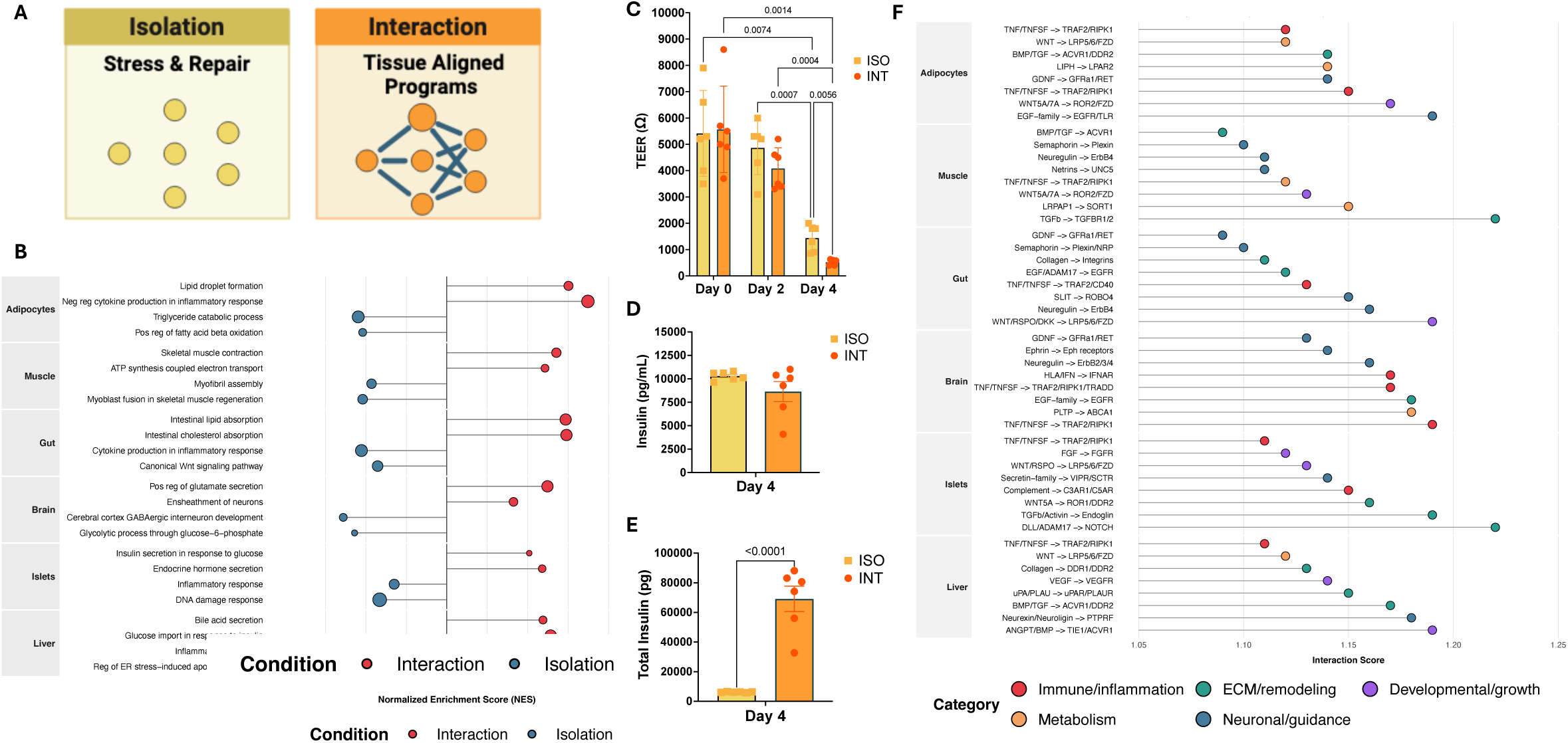
Multi-tissue interaction redirects engineered tissue state toward compartment-aligned functional programs. **(A)** Summary schematic highlighting the major shared perfusion-associated tissue-state changes observed under the Mid nutrient condition, 12 mM glucose/2% Lipid Mixture 1. **(B)** GSEA comparing interaction versus isolation across six tissues under the Mid nutrient condition. Normalized enrichment scores (NES) are shown for selected GO Biological Process pathways; positive NES indicates enrichment in interaction and negative NES indicates enrichment in isolation. Dot size reflects statistical significance as −log₁₀ P value. Two representative pathways are shown per tissue. RNA-seq data: n = 3 per condition per tissue from two independent experiments. **(C)** TEER of gut epithelium over time in isolation (ISO) and interaction (INT) conditions. Data represent mean ± SEM; statistical significance was determined by two-way repeated-measures ANOVA with condition and time as factors, followed by Tukey’s multiple-comparisons test. n = 6 per condition from two independent experiments. **(D)** Insulin concentration in media and **(E)** total insulin secretion under the Mid nutrient condition. Total insulin secretion was calculated as insulin concentration × total media volume to account for culture-volume differences between isolation and interaction conditions. Data represent mean ± SEM; statistical significance was determined by unpaired t-test. n = 6 per condition from two independent experiments. **(F)** Selected tissue-resolved ligand–receptor signatures inferred from RNA-seq profiles using BulkSignalR with the default ligand–receptor database. Displayed pairs were selected based on statistical support, inferred interaction score, and biological relevance to the interaction-associated tissue-state programs shown in (B). Tissue labels indicate the compartment in which each interaction score was calculated; pairs were not required to be exclusive to a single tissue and may be detected across compartments at different inferred levels. Colors indicate functional signaling category: immune/inflammation, red; metabolism, orange; ECM/remodeling, teal; neuronal/guidance, blue; developmental/growth, purple. Interaction score reflects the BulkSignalR-inferred ligand–receptor score under the applied thresholds.

Shared perfusion produced tissue-specific transcriptional changes rather than a uniform response across compartments (Fig.2B; Fig.S2). In adipocytes, interaction enriched lipid droplet formation and negative regulation of cytokine production in inflammatory response, whereas isolation enriched triglyceride catabolism and positive regulation of fatty acid beta-oxidation. In skeletal muscle, interaction enriched skeletal muscle contraction and ATP synthesis-coupled electron transport, while isolation enriched myofibril assembly and myoblast fusion involved in skeletal muscle regeneration. These patterns suggest that shared perfusion shifted adipocytes toward lipid storage and inflammatory restraint and skeletal muscle toward contractile bioenergetic programs, while isolation favored adipocyte catabolism and muscle repair-associated programs.

Interaction also remodeled neural, endocrine, hepatic, and epithelial states (Fig.2B; Fig.S2). Brain organoids shifted toward glutamate secretion and ensheathment of neurons under interaction, whereas isolation enriched GABAergic interneuron development and glycolytic process through glucose-6-phosphate. Pancreatic islets enriched insulin secretion in response to glucose and endocrine hormone secretion under interaction, while isolation enriched inflammatory response and DNA damage response. Liver organoids enriched bile acid secretion and glucose import in response to insulin under interaction, whereas isolation enriched inflammatory response and ER stress-induced apoptotic signaling. In gut epithelium, interaction enriched intestinal lipid and cholesterol absorption, whereas isolation enriched canonical Wnt signaling and inflammatory cytokine production, consistent with stronger renewal-associated and surveillance programs in isolated cultures. TEER declined over time in both conditions and was lower in interaction by day 4 (Fig.2C). This functional change occurred alongside interaction-associated absorptive programs and isolation-associated Wnt signaling, suggesting a renewal-to-absorptive shift in the present shared-media format, consistent with links among Wnt signaling, intestinal epithelial renewal, differentiation state, and barrier-function readouts in engineered intestinal models ^38–41^.

Islet responses provided a second functional example. Insulin concentration in media was not significantly different between isolation and interaction under the Mid nutrient condition (Fig.2D). However, because the interacting MPS contains a larger shared perfusate volume, concentration alone does not capture total endocrine output ^42^. After accounting for media volume, interacting islets produced significantly more total insulin than isolated islets (Fig.2E). This increase was concordant with enrichment of insulin secretion in response to glucose and endocrine hormone secretion, supporting a more endocrine-functional islet state under shared perfusion.

Ligand–receptor inference further supported a tissue-resolved communication landscape in the interacting circuit (Fig.2F). Ligand–receptor pairs were inferred from tissue-resolved RNA-seq profiles; tissue labels indicate the compartment in which the interaction score was calculated, and the resulting network reflects putative signaling relationships across the interacting circuit. We therefore visualized selected pairs that were statistically supported, detected across tissue compartments at varying inferred levels, and biologically aligned with the interaction-associated tissue-state programs. These pairs included immune/inflammatory signals such as *TNF*/*TNFSF*–*TRAF2*/*RIPK1*; metabolic and trophic pathways including *WNT*/*RSPO*–*LRP5/6/FZD* and EGF-family–*EGFR/*ErbB signaling; ECM/remodeling-associated *BMP/TGF*–*ACVR/DDR* interactions; and neural/guidance-associated semaphorin–plexin, neuregulin–ErbB, *GDNF*–*GFRA/RET*, netrin–*UNC5*, and ephrin receptor signaling. These inferred signaling classes paralleled the interaction-associated tissue-state changes observed by pathway analysis.

Collectively, these data indicate that multi-tissue interaction redirected engineered tissue state in a compartment-specific manner. Isolation preferentially retained maintenance, stress, repair, inflammatory, or renewal-associated programs, while shared perfusion promoted tissue-aligned states including adipocyte lipid storage, muscle contractile bioenergetics, gut absorptive metabolism, brain organoid neural-support programs, islet endocrine secretion, and liver organoid insulin-responsive glucose and bile-acid-associated programs. Functional endpoints captured only part of this remodeling: gut absorptive programs coincided with reduced TEER, while islet endocrine programming coincided with increased total insulin output. These findings established the shared-perfusion reference state for subsequent nutrient and drug perturbation studies.

### Increased nutrient availability shifts the six-tissue MPS from maintenance toward compensatory metabolic remodeling

We next asked whether the interacting six-tissue MPS could resolve graded nutrient availability after assembly into shared perfusion. Six-tissue circuits were exposed to Low nutrient conditions, defined as 6 mM glucose with 0.5% Lipid Mixture 1, or Mid nutrient conditions, defined as 12 mM glucose with 2% Lipid Mixture 1 (Fig.3A). Circulating media were collected after two and four days for functional, endocrine, inflammatory, and metabolomic profiling, and tissues were collected on day 4 for transcriptomic analysis.

**Figure 3.**
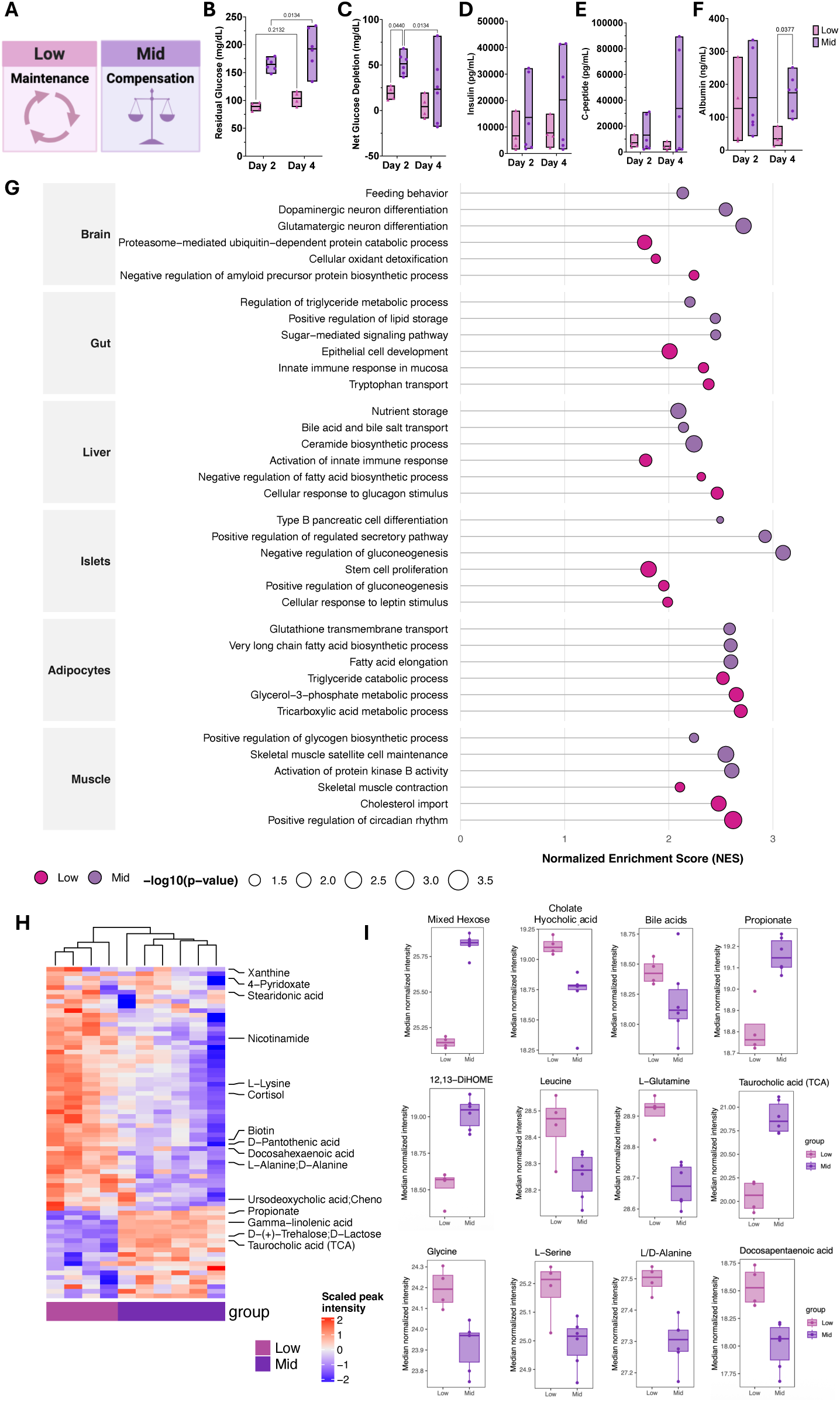
Increased nutrient availability shifts the six-tissue MPS from maintenance toward compensatory metabolic remodeling. **(A)** Summary schematic highlighting the major circuit-level changes observed between Low nutrient conditions, 6 mM glucose/0.5% Lipid Mixture 1, and Mid nutrient conditions, 12 mM glucose/2% Lipid Mixture 1. Circulating media were collected after two and four days of shared perfusion for functional, endocrine, inflammatory, and metabolomic profiling. **(B–F)** Circulating media and functional readouts from Low and Mid conditions: **(B)** residual glucose concentration, **(C)** net glucose depletion, **(D)** insulin concentration, **(E)** C-peptide concentration, and **(F)** albumin concentration. Net glucose depletion was calculated by subtracting residual glucose concentration from starting glucose concentration. Data are shown as box plots with center line indicating mean and bounds indicating min–max. Statistical significance was determined by two-way repeated-measures ANOVA with nutrient condition and time as factors, followed by Tukey’s multiple-comparisons test. n = 4 for Low and n = 6 for Mid from two independent experiments. **(G)** GSEA comparing Low vs Mid conditions across six tissues. Normalized enrichment scores (NES) are shown for selected GO Biological Process pathways. Dot size reflects statistical significance as −log₁₀ P value. Three representative pathways are shown per tissue. **(H)** Heatmap of significantly differentially abundant metabolites detected in circulating media from Low versus Mid conditions. Values are shown as scaled peak intensity across individual samples. **(I)** Normalized median intensity of selected metabolites and metabolite groups altered under Mid nutrient conditions, n = 4 for Low and n = 6 for Mid from two independent experiments.

The Mid nutrient condition produced a circuit-level response consistent with compensatory nutrient adaptation. Residual glucose was significantly higher under Mid conditions by day 4, whereas net glucose depletion decreased from day 2 to day 4 despite greater starting nutrient availability (Fig.3B,C). In parallel, insulin, C-peptide, and albumin were elevated under Mid conditions, with C-peptide increasing from day 2 to day 4 (Fig.3D-F). Additional shared-media readouts detected adiponectin and IL-8, while TEER did not differ significantly between Low and Mid conditions within the interacting circuit (Fig.S3A–C). Thus, the Low-to-Mid transition increased endocrine and liver organoid-associated secretory output and remodeled soluble signaling in the shared perfusate without producing a measurable nutrient-dependent divergence in epithelial barrier resistance.

Transcriptomic profiling revealed that this functional shift was accompanied by tissue-specific nutrient-state remodeling (Fig.3G; Fig.S3D,E). Under Low conditions, the circuit retained tissue-specific programs consistent with lower-nutrient maintenance, surveillance, and substrate mobilization (Fig. 3G; Fig. S3D,E). Adipocytes enriched triglyceride catabolism, glycerol-3-phosphate metabolism, and tricarboxylic acid metabolism. Liver organoids enriched cellular response to glucagon stimulus and negative regulation of fatty acid biosynthesis. Skeletal muscle enriched muscle contraction, cholesterol import, and positive regulation of circadian rhythm. Low conditions also enriched gut epithelial development, mucosal innate immune response, and tryptophan transport, alongside brain organoid programs related to proteasome-mediated protein catabolism, cellular oxidant detoxification, and negative regulation of amyloid precursor protein biosynthesis.

Mid conditions shifted the interacting circuit toward anabolic, secretory, storage, and nutrient-handling programs (Fig.3G; Fig.S3D,E). Islets enriched type B pancreatic cell differentiation and positive regulation of regulated secretory pathway, consistent with increased insulin and C-peptide accumulation and engagement of β-cell secretory machinery ^43^. Liver organoids enriched nutrient storage, bile acid and bile salt transport, and ceramide biosynthetic process, indicating increased nutrient handling, lipid/sphingolipid remodeling, and bile-acid-associated signaling^44,45^. Skeletal muscle enriched biosynthesis and AKT-associated signaling, consistent with insulin-linked anabolic and nutrient-handling pathways ^46^. Adipocytes enriched very long-chain fatty acid biosynthesis, fatty acid elongation, *HIF1α* signaling, and glutathione transmembrane transport, suggesting lipid biosynthetic remodeling alongside hypoxia-and redox-associated adaptation ^47^. Gut epithelium enriched regulation of triglyceride metabolism, lipid storage, and sugar-mediated signaling, while brain organoids enriched feeding-associated gene programs, dopaminergic neuron differentiation, and glutamatergic differentiation programs, consistent with nutrient-sensitive gut and neural transcriptional responses involved in metabolic-state sensing ^48,49^.

The circulating metabolome further showed that the Low-to-Mid transition did not simply increase all nutrient-associated features. Low and Mid samples separated by global metabolite profiles, indicating remodeling of the shared perfusate with increased glucose/lipid availability (Fig.3H,I). Mid conditions increased mixed hexoses, propionate, 12,13-DiHOME, and taurocholic acid, while decreasing leucine, glutamine, glycine, serine, alanine, docosapentaenoic acid, cholate/hyocholic acid, and a multi-annotated bile-acid feature (Fig.3I). The combination of elevated carbohydrate-associated features with lower amino acids and altered bile-acid/lipid-derived metabolites suggests coordinated changes in nutrient utilization, secretion, and exchange across the interacting six-tissue circuit.

Collectively, the Low-to-Mid comparison indicates that increased nutrient availability drove a compensated metabolic state in the interacting six-tissue MPS. Under Mid conditions, increased insulin, C-peptide, albumin, anabolic transcriptional programs, and metabolite remodeling showed that the circuit detected and responded to increased glucose/lipid exposure. The concurrent decline in net glucose depletion over time indicates that this adaptation was not proportional to nutrient availability, establishing a reference point for testing whether further nutrient escalation shifts the system toward stress-associated dysfunction.

### Sustained nutrient overload shifts the six-tissue MPS toward stress-associated metabolic dysfunction

We next compared Mid and High nutrient conditions to determine how the interacting six-tissue MPS responds to sustained nutrient overload. Six-tissue circuits were exposed to Mid nutrient conditions, defined as 12 mM glucose with 2% Lipid Mixture 1, or High nutrient conditions, defined as 18 mM glucose with 4% Lipid Mixture 1, with High conditions designed to model selected features of sustained nutrient excess (Fig.4A). Relative to Mid, High conditions increased residual glucose over time, while net glucose depletion declined from day 2 to day 4 (Fig.4B,C). C-peptide showed a modest nutrient-associated increase, although the difference between Mid and High was not significant (Fig.4D), suggesting that endocrine output did not scale proportionally with the greater nutrient load. Glucagon increased over time in both Mid and High conditions and was higher under High conditions by day 4 (Fig.4E). Under sustained nutrient availability, elevated glucagon suggests that the circuit did not fully suppress counter-regulatory endocrine signaling, a pattern consistent with clinical studies linking hyperglucagonemia to impaired glucose regulation and altered liver–α-cell communication ^50–52^. Together, increased residual glucose, declining net glucose depletion, incomplete C-peptide scaling, and elevated glucagon indicate that High nutrient exposure reduced apparent glucose-handling efficiency and altered counter-regulatory endocrine signaling.

**Figure 4.**
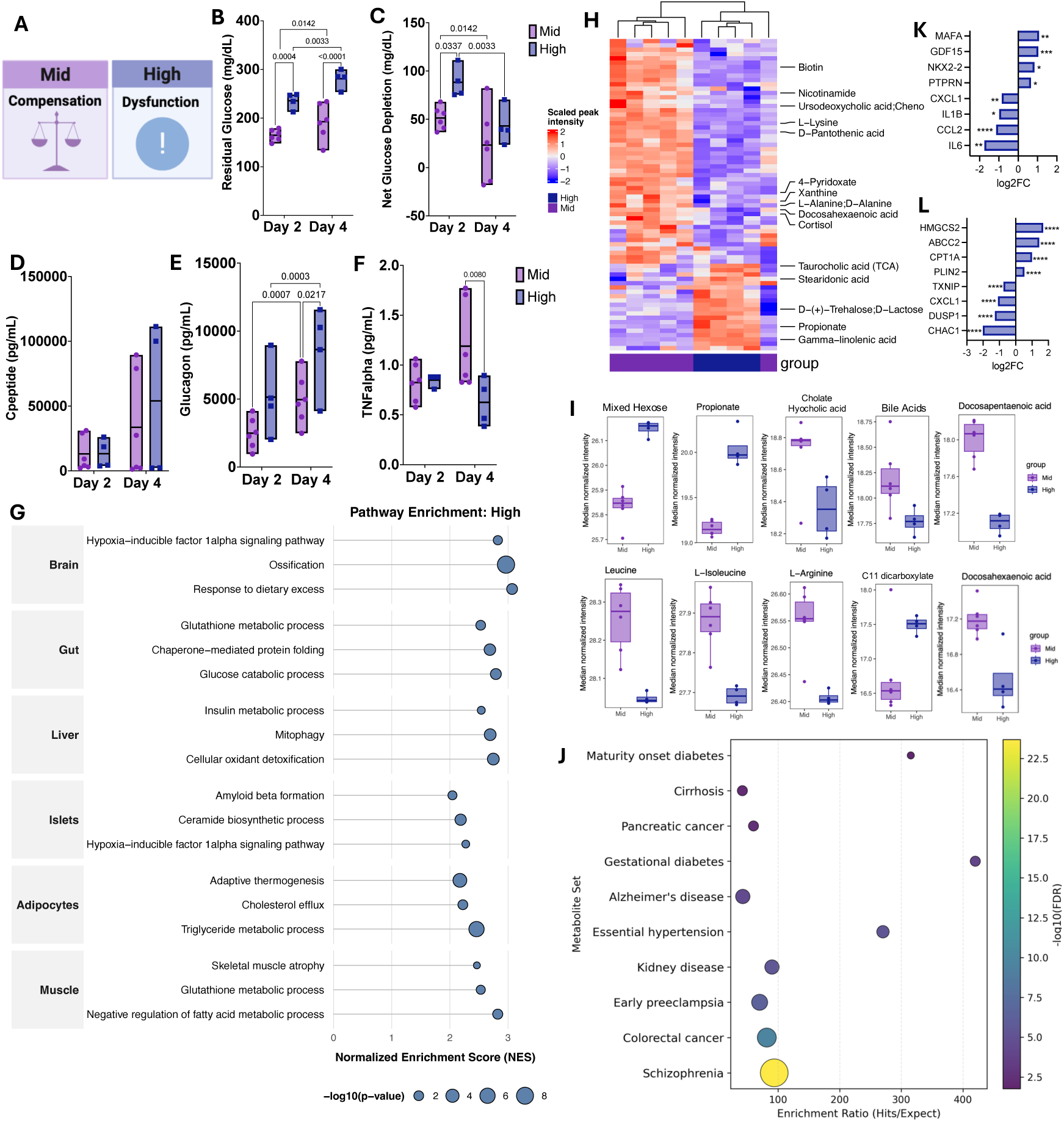
Sustained nutrient overload shifts the interacting six-tissue MPS toward stress-associated metabolic dysfunction. **(A)** Summary schematic highlighting the major circuit-level changes observed under Mid nutrient conditions, 12 mM glucose/2% Lipid Mixture 1, or High nutrient conditions, 18 mM glucose/4% Lipid Mixture 1. Circulating media were collected after two and four days of shared perfusion for functional, endocrine, inflammatory, and metabolomic profiling; tissues were collected on day 4 for transcriptomic analysis. **(B–F)** Circulating media and functional readouts from Mid and High nutrient conditions: **(B)** residual glucose concentration, **(C)** net glucose depletion, **(D)** C-peptide concentration, **(E)** glucagon concentration, and **(F)** TNFα concentration. Net glucose depletion was calculated by subtracting residual glucose concentration from starting glucose concentration. Data are shown as box plots with center line indicating mean and bounds indicating min–max. Statistical significance was determined by two-way repeated-measures ANOVA with nutrient condition and time as factors, followed by Tukey’s multiple-comparisons test. n = 6 for Mid and n = 4 for High from two independent experiments. **(G)** GSEA comparing High versus Mid nutrient conditions across six tissues. Normalized enrichment scores (NES) are shown for selected GO Biological Process pathways. Dot size reflects statistical significance as −log₁₀ P value. Three selected pathways are shown per tissue. RNA-seq data: n = 4 per condition per tissue from two independent experiments. **(H)** Heatmap of significantly differentially abundant metabolite features detected in circulating media from Mid and High nutrient conditions. Values are shown as scaled peak intensity across individual samples. **(I)** Normalized median intensity of selected metabolite features and metabolite groups altered under High nutrient conditions. n = 6 for Mid and n = 4 for High from two independent experiments. **(J)** HMDB metabolite-set enrichment analysis of significantly differentially abundant metabolite features in circulating media comparing High versus Mid nutrient conditions. Metabolite sets were considered significant at FDR < 0.05. Enrichments indicate overlap with curated disease-associated reference metabolite sets. **(K)** Pancreatic islet and **(L)** liver organoid transcriptional responses to High nutrient exposure comparing isolation and interaction conditions. Bars show RNA-seq-derived log₂ fold change for selected genes; positive values indicate higher expression in isolation and negative values indicate higher expression in interaction. Differential expression was calculated using DESeq2 with Benjamini-Hochberg correction. n = 3 per condition for each tissue module. Significance indicates Benjamini-Hochberg-adjusted P values; *P < 0.05, **P < 0.01, ***P < 0.001, and ****P < 0.0001.

Transcriptomic profiling suggested that High conditions engaged stress and quality-control programs across metabolic hub tissues (Fig.4G; Fig.S4). Liver organoids enriched ER-associated degradation, unfolded protein response, macroautophagy, mitophagy, oxidant detoxification, proteasome-mediated protein catabolism, ferroptosis-associated programs, and reduced cholesterol biosynthetic regulation. These signatures are consistent with proteostatic, oxidative, and lipid-redox stress remodeling under lipid-rich conditions ^53,54^. Liver organoid-associated stress programs coincided with extracellular metabolite remodeling, including altered bile-acid features, increased taurocholic acid and C11 dicarboxylate, and reduced DPA/DHA, suggesting disruption of lipid-, bile-acid-, and oxidative metabolism-associated features under High conditions (Fig.4H,I).

Pancreatic islets also showed stress-associated remodeling under High conditions, with enrichment of ceramide and sphingolipid biosynthesis, amyloid beta formation, macrophage activation programs, HIF-1α signaling, DNA repair, and triglyceride-rich lipoprotein particle remodeling (Fig.4G; Fig.S4). These signatures are consistent with lipid-, inflammatory-, and proteotoxic stress-associated programs implicated in β-cell dysfunction in type 2 diabetes ^55,56^. Functionally, C-peptide did not significantly increase from Mid to High despite greater nutrient burden, indicating limited scaling of islet secretory output under High conditions (Fig.4D).

High nutrient exposure also altered insulin-sensitive tissue compartments. Skeletal muscle enriched atrophy, starvation-response, inflammatory, glutathione metabolic, lipid-oxidation, ketone-regulatory, and fatty-acid metabolic programs despite extracellular nutrient abundance (Fig.4G; Fig.S4), consistent with catabolic and stress-responsive remodeling linked to impaired nutrient handling in insulin-resistant states ^57^. Adipocytes enriched adaptive thermogenesis, brown fat cell differentiation, cholesterol efflux, triglyceride metabolism, fatty acid beta-oxidation, D-glucose import, insulin response, plasma lipoprotein particle clearance, and macroautophagy (Fig.4G; Fig.S4). These programs suggest concurrent metabolic adaptation and quality-control remodeling under sustained High nutrient exposure, consistent with the context-dependent role of brown/beige fat programs in energy dissipation ^58^.

Gut epithelium and brain organoids also responded to the High nutrient shared perfusate. Gut epithelium enriched redox, oxidative phosphorylation, protein-folding, misfolded protein response, proteasomal catabolism, stress-activated kinase signaling, inflammatory cytokine production, macroautophagy, and glucose catabolic programs (Fig.4G; Fig.S4). These pathways suggest redox buffering, protein quality-control remodeling, inflammatory signaling, and energetic stress within the intestinal compartment, consistent with links between intestinal barrier-associated inflammatory signaling and metabolic dysfunction ^59^. Brain organoids enriched *HIF-1α* signaling, excess nutrient response-associated programs, *TGFβ/BMP*-associated pathways, collagen fibril organization, and cellular stress-response programs (Fig.4G; Fig.S4). The brain organoid compartment is coupled to the circuit through shared perfusate, without vascular, autonomic, or behavioral inputs, linking these transcriptional signatures to nutrient-sensitive humoral cues.

Conditioned-media metabolomics further supported a High-associated shift in the shared extracellular environment (Fig.4H-J). Relative to Mid, High conditions generated a distinct extracellular metabolite profile characterized by increased mixed hexoses, propionate, taurocholic acid, and C11 dicarboxylate, together with reduced cholate/hyocholic acid, total bile-acid signal, L-arginine, D-pantothenic acid, and DPA/DHA (Fig.4I). These changes suggest altered substrate handling and bile-acid composition, consistent with the role of bile-acid signaling in metabolic regulation and metabolic disease ^60,61^. Decreased D-pantothenic acid is consistent with altered pantothenate/CoA-linked metabolic capacity and broader shifts in energy and fatty-acid metabolism ^62^. Leucine and isoleucine were also decreased under High conditions, consistent with altered branched-chain amino acid (BCAA) handling; dysregulated BCAA metabolism has been linked to insulin resistance and metabolic disease, diabetes-related complications, and neuropsychiatric disease-associated metabolic disruption ^63–65^. Metabolite set enrichment analysis further showed that the High-state extracellular profile overlapped with reference metabolite sets associated with chronic metabolic, vascular, inflammatory, and neuroendocrine disease contexts, including kidney disease, hypertension, gestational diabetes, cirrhosis, Alzheimer’s disease, and maturity-onset diabetes (Fig.4J). Together, these metabolomic changes indicate that sustained nutrient overload shifted the shared perfusate toward biochemical patterns associated with chronic tissue stress and metabolic dysfunction.

Finally, we asked whether multi-tissue interaction modified High nutrient responses relative to isolation. In both islets and liver organoids, interaction shifted High nutrient responses toward inflammatory and nutrient-stress-associated gene expression relative to isolated culture (Fig.4K,L). In islets, interaction increased inflammatory and chemokine-associated genes, including *IL6, IL1B, CCL2*, and *CXCL1*, whereas isolation showed higher expression of genes associated with β-cell endocrine identity and insulin secretory function, including *MAFA, NKX2-2,* and *PTPRN*, as well as stress-responsive endocrine adaptation marked by GDF15 (Fig.4K). In liver organoids, interaction increased genes associated with amino acid/ER-integrated stress signaling, stress-responsive *MAPK* regulation, inflammatory chemokine activation, and glucose/nutrient stress sensing, including *CHAC1, DUSP1, CXCL1*, and *TXNIP*. By contrast, isolation showed higher expression of genes associated with lipid-droplet handling, mitochondrial fatty-acid oxidation, efflux/transport function, and ketone/cholesterol-associated metabolic adaptation, including *PLIN2, CPT1A, ABCC2*, and *HMGCS2* (Fig.4L). Thus, multi-tissue context supported compartment-aligned programs under the Mid reference condition and amplified inflammatory and nutrient-stress cues in key glucose-regulatory compartments under High nutrient exposure.

Collectively, these data indicate that High nutrient exposure shifted the six-tissue MPS from a compensatory nutrient-adapted state toward stress-associated metabolic dysfunction. This state was marked by increased residual glucose, reduced net glucose depletion over time, incomplete scaling of endocrine output, altered glucagon regulation, tissue-specific stress and quality-control programs, and remodeling of the shared extracellular metabolite environment. High nutrient exposure also changed the effect of tissue interaction itself: shared perfusion supported compartment-aligned programs under Mid conditions and amplified stress-associated programs in liver organoid and islet compartments under High conditions.

### Metformin and semaglutide produce distinct response modes under High nutrient conditions

To determine whether established anti-diabetic therapies could modify circuit-level responses under sustained nutrient overload, six-tissue MPS maintained under High nutrient conditions were treated with metformin or semaglutide during shared perfusion (Fig.5A). Media collected on days 2 and 4 were used to assess net glucose depletion, endocrine output, inflammatory signaling, epithelial barrier resistance, extracellular metabolites, and tissue transcriptional responses. Drug responses were interpreted as circuit-level tissue and media-state changes, recognizing that the platform does not capture full organismal pharmacology, renal clearance, appetite regulation, body-weight-associated effects, or long-term exposure kinetics.

**Figure 5.**
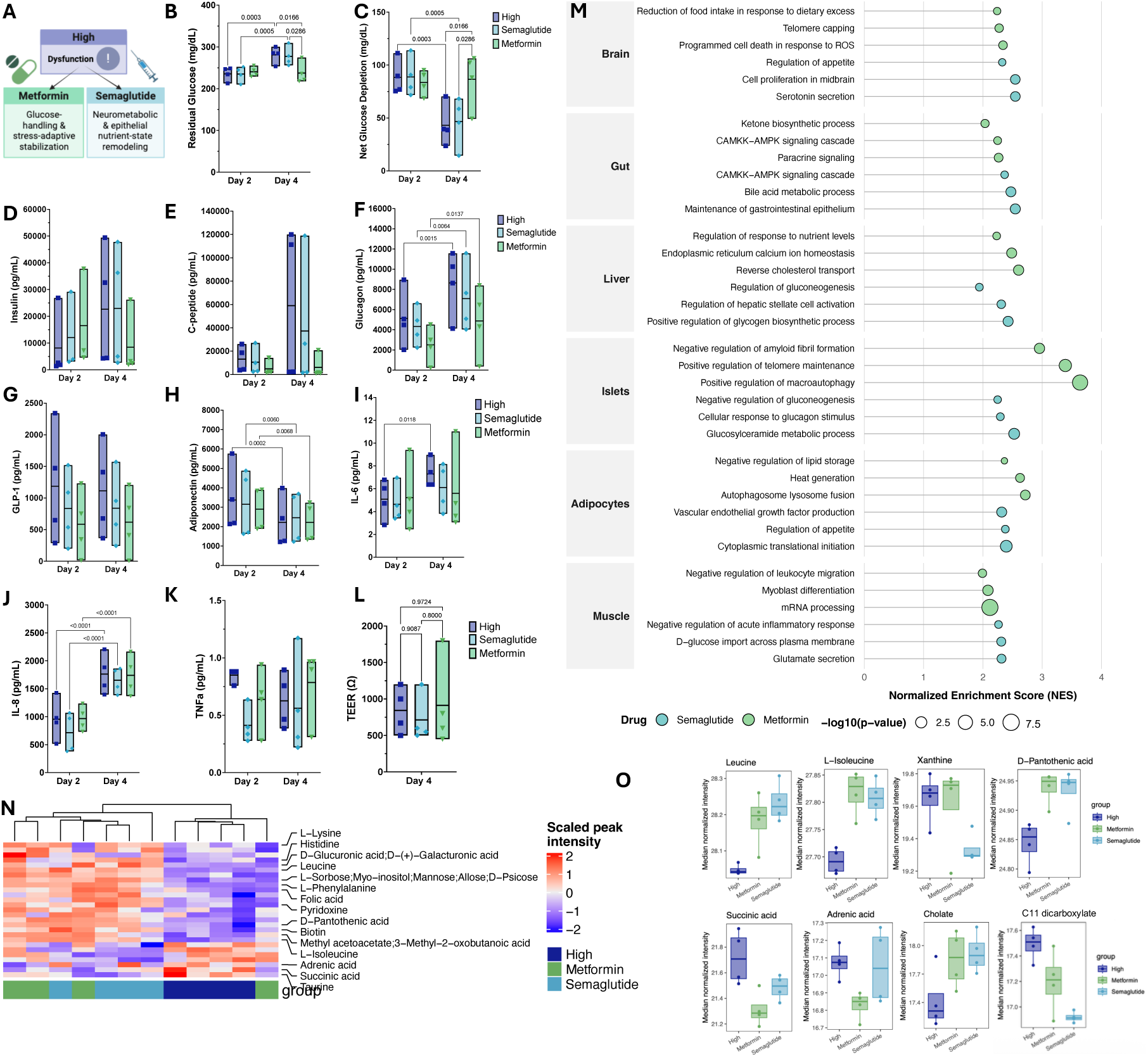
Metformin and semaglutide induce distinct functional and transcriptional responses under High nutrient conditions. **(A)** Summary schematic highlighting the major circuit-level changes observed under High nutrient conditions, 18 mM glucose/4% Lipid Mixture 1, with or without metformin or semaglutide. Drugs were administered on days 0 and 2 during shared perfusion, and circulating media were collected after two and four days for functional, endocrine, inflammatory, barrier, and metabolomic profiling. **(B–K)** Circulating media readouts from High, High + metformin, and High + semaglutide conditions: **(B)** residual glucose concentration, **(C)** net glucose depletion, **(D)** insulin concentration, **(E)** C-peptide concentration, **(F)** glucagon concentration, **(G)** GLP-1 concentration, **(H)** adiponectin concentration, **(I)** IL-6 concentration, **(J)** IL-8 concentration, and **(K)** TNFα concentration. Net glucose depletion was calculated by subtracting residual glucose concentration from starting glucose concentration. Data are shown as box plots with center line indicating mean and bounds indicating min–max. Statistical significance was determined by two-way repeated-measures ANOVA with treatment condition and time as factors, followed by Tukey’s multiple-comparisons test. n = 4 per condition from two independent experiments. **(L)** TEER of the gut monolayer under High, High + metformin, and High + semaglutide conditions. Data are shown as box plots with center line indicating mean and bounds indicating min–max. Statistical significance was determined by unpaired t-tests. n = 4-6 from two independent experiments. **(M)** GSEA comparing drug-treated conditions versus untreated High conditions across six tissues. Normalized enrichment scores (NES) are shown for selected GO Biological Process pathways. Color indicates drug treatment: metformin, green; semaglutide, light blue. Dot size reflects statistical significance as −log₁₀ P value. Three representative pathways are shown per tissue. **(N)** Heatmap of significantly differentially abundant metabolites detected in circulating media from High, High + metformin, and High + semaglutide conditions. Values are shown as scaled peak intensity across individual samples. **(O)** Normalized median intensity of selected metabolites and metabolite groups altered by drug treatment under High nutrient conditions.

Despite equivalent glucose input across groups, treatment effects on shared-media glucose handling diverged over time. Residual glucose increased from day 2 to day 4 in untreated High and semaglutide-treated MPS, while metformin-treated MPS maintained lower residual glucose by day 4 and showed increased net glucose depletion relative to both groups (Fig.5B,C). Consistent with metformin biology involving hepatic glucose regulation and intestinal epithelial mitochondrial metabolism, this improvement occurred without a proportional increase in insulin or C-peptide accumulation, supporting improved shared-media glucose handling without a measurable increase in β-cell secretory burden over this timeframe (Fig.5D,E) ^66–68^.

Other circulating readouts indicated partial and endpoint-specific drug effects across the four-day exposure window. Glucagon increased from day 2 to day 4 across conditions, although the metformin-treated group appeared lower than untreated High and semaglutide-treated MPS at day 4, consistent with partial attenuation of counter-regulatory endocrine signaling (Fig.5F) ^51,52^. GLP-1 did not show a clear treatment-associated increase, consistent with semaglutide acting through receptor agonism without increasing endogenous GLP-1 abundance in the shared perfusate (Fig.5G).

Adiponectin decreased over time across groups, IL-6 increased in untreated High but was comparatively restrained in both drug-treated groups, IL-8 increased over time across conditions, and TNFα remained variable without a clear treatment-specific pattern (Fig.5H-K). TEER was not significantly different between groups at day 4 (Fig.5L). Thus, metformin produced the clearest functional rescue at the level of shared-media glucose handling.

At the transcriptional level, the two drugs induced separable tissue-specific programs under High nutrient conditions (Fig.5M). Semaglutide preferentially remodeled gut epithelium, liver organoid, islet, and brain organoid transcriptional programs linked to nutrient sensing, epithelial maintenance, endocrine response, and neurometabolic state. In gut epithelium, semaglutide enriched bile acid metabolic process, maintenance of gastrointestinal epithelium, *CAMKK–AMPK* signaling, and paracrine signaling, suggesting engagement of epithelial nutrient sensing, barrier-associated maintenance, and local secretory communication programs (Fig.5M; Fig.S5A–C). In liver organoids, semaglutide enriched positive regulation of glycogen biosynthetic process, regulation of gluconeogenesis, regulation of hepatic stellate cell activation, and regulation of response to nutrient levels, consistent with carbohydrate-handling and remodeling-associated programs under High nutrient conditions (Fig.5M; Fig.S5A–C). This is broadly aligned with reported effects of semaglutide on metabolic liver disease and fibrosis-associated endpoints ^69,70^.

Semaglutide-associated pathway changes extended across endocrine, peripheral metabolic, and neural compartments, but these transcriptional effects were not matched by an acute improvement in net glucose depletion. In islets, semaglutide enriched cellular response to glucagon stimulus, negative regulation of gluconeogenesis, and glucosylceramide metabolic process, indicating remodeling of endocrine stimulus-response and metabolic stress-associated pathways (Fig.5M; Fig.S5A–C). These programs are consistent with GLP-1-linked regulation of islet stimulus-secretion biology, although net glucose depletion did not increase over the measured window (Fig.5B–F)^71^. In skeletal muscle, semaglutide enriched D-glucose import across the plasma membrane, glutamate secretion, and negative regulation of acute inflammatory response, suggesting partial engagement of peripheral glucose-handling and anti-inflammatory programs (Fig.5M; Fig.S5A–C). In adipocytes, semaglutide-associated terms included vascular endothelial growth factor production, appetite-associated transcriptional programs, and cytoplasmic translational initiation, consistent with secretory, vascular-associated, and biosynthetic remodeling (Fig.5M; Fig.S5A–C).

Brain organoids showed a prominent semaglutide-associated transcriptional response, with enrichment of appetite-associated transcriptional programs, serotonin secretion, cell proliferation in midbrain, and, in the broader supplemental pathway view, neuronal morphogenesis and synaptic transmission-associated programs (Fig.5M; Fig.S5A–C). This profile is consistent with GLP-1-associated neurometabolic biology, including clinical evidence that semaglutide can reduce appetite, energy intake, and food cravings, and alter brain responses to food-related or taste-associated stimuli ^72–75^. In this reduced circuit, the brain organoid compartment is coupled through shared perfusate, with vascular, autonomic, or behavioral inputs being absent, linking these transcriptional responses to drug- and nutrient-sensitive humoral cues. Together, the semaglutide response indicates broad tissue-state remodeling across neural, epithelial, liver organoid, endocrine, and peripheral metabolic compartments, with limited short-term impact on aggregate glucose depletion.

Metformin produced a distinct stabilization-associated program more closely aligned with the functional glucose-handling phenotype (Fig.5M; Fig.S5B,C). In liver organoids, metformin enriched reverse cholesterol transport, endoplasmic reticulum calcium ion homeostasis, and regulation of response to nutrient levels, consistent with lipid transport, ER homeostasis, and nutrient-stress adaptation under High nutrient conditions (Fig.5M; Fig.S5C). These pathways align with known metformin biology involving hepatic metabolic regulation, redox control, and altered glucose production, and were concordant with the functional preservation of net glucose depletion (Fig.5B,C) ^67,76,77^. These transcriptional signatures support partial liver organoid stabilization within the High nutrient state.

In islets, metformin enriched negative regulation of amyloid fibril formation, positive regulation of macroautophagy, positive regulation of telomere maintenance, and cellular response to calcium ion, consistent with proteostatic and stress-adaptive remodeling (Fig.5M; Fig.S5C). High nutrient exposure enriched amyloid fibril formation-associated programs, whereas metformin shifted the pathway direction toward negative regulation of amyloid fibril formation, suggesting partial transcriptional counter-regulation of a proteotoxic β-cell stress-associated feature. Islet amyloid polypeptide aggregation is a well-established feature of type 2 diabetes-associated β-cell dysfunction, and autophagy/lysosomal degradation has been shown to protect β cells from human IAPP-induced proteotoxicity ^78–80^. This pattern is compatible with pathway-level proteostatic remodeling in nutrient-stressed islets ^56,81^. In adipocytes, metformin enriched negative regulation of lipid storage, heat generation, and autophagosome–lysosome fusion, indicating altered lipid handling, energy expenditure-associated programming, and autophagic remodeling (Fig.5M; Fig.S5C). In skeletal muscle, metformin enriched myoblast differentiation, mRNA processing, and negative regulation of leukocyte migration, with cAMP–PKA and WNT-associated signaling terms (Fig.5M; Fig.S5C). Across these compartments, metformin preferentially engaged proteostatic, autophagic, lipid-remodeling, and peripheral metabolic-flexibility programs.

The brain organoid compartment further separated the two drug responses. Semaglutide shifted High nutrient-exposed brain organoids toward appetite-associated, serotonergic, midbrain, morphogenesis, and synaptic transcriptional programs, whereas metformin enriched telomere capping, programmed cell death in response to reactive oxygen species, reduction of food intake in response to dietary excess, interferon-mediated signaling, leukocyte migration involved in inflammatory response, programmed cell death in response to ROS, and synaptic depression-associated pathways (Fig.5M; Fig.S5C). This profile suggests a distinct stress-adaptive transcriptional response, consistent with reported links between metformin, CNS-relevant AMPK signaling, autophagy, and stress-response pathways ^82^. Thus, the brain organoid response reinforced the broader separation between semaglutide-associated neurometabolic remodeling and metformin-associated stress-adaptive remodeling under High nutrient exposure.

The metabolomics data supported overlapping but distinct biochemical effects of treatment in the shared perfusate. Both drugs shifted the extracellular metabolite profile relative to untreated High conditions, indicating broad biochemical remodeling in the shared circulating media environment (Fig.5N). Selected metabolite features showed increased branched-chain amino acids, including leucine and isoleucine, with treatment, while semaglutide reduced xanthine and C11 dicarboxylate relative to untreated High (Fig.5O). Metformin increased pantothenic acid and cholate and reduced succinic acid relative to untreated High, consistent with altered substrate handling, mitochondrial/redox-associated metabolism, and bile-acid-linked metabolic communication (Fig.5O) ^45,61,77^. These changes report altered metabolite availability or handling in the circulating compartment, not tissue-specific metabolic flux.

Overall, semaglutide and metformin produced distinct response modes under High nutrient conditions. Metformin most effectively preserved circuit-level net glucose depletion and induced a liver organoid-, islet-, adipocyte-, and muscle-associated stabilization program involving nutrient response, reverse cholesterol transport, ER homeostasis, macroautophagy, lipid storage regulation, and glucose handling.

Semaglutide produced a broader gut–brain organoid–islet–liver organoid remodeling signature involving epithelial maintenance, bile-acid metabolism, appetite- and serotonin-associated brain organoid programs, glycogen biosynthesis, and endocrine stimulus-response programs, with more limited acute improvement in net glucose depletion. Thus, the interacting six-tissue MPS separated two therapeutic response modes under sustained nutrient overload: metformin preferentially preserved shared-media glucose handling with stabilization-associated tissue programs, whereas semaglutide primarily reshaped nutrient-sensing, epithelial, endocrine, and neurometabolic transcriptional states.

### Discussion and Conclusion

This study establishes a perfused human six-tissue MPS, the AnthroHive and MOTIVE-6, for investigating how shared circulation, nutrient availability, and metabolic therapies shape glucose-regulatory tissue states.

Existing MPS models have reproduced important aspects of metabolic crosstalk across selected tissue axes, including adipose–liver, pancreas–muscle–liver, liver–islet, gut–liver, and gut–liver–brain systems ^23–29,36,37,83–85^. Glucose regulation spans a broader network of absorptive, endocrine, hepatic, storage, contractile, inflammatory, and neural-associated compartments whose coordinated signaling contributes to nutrient sensing, hormone output, substrate partitioning, inflammatory tone, and systemic metabolic balance ^11,46,86^. The six-tissue MPS developed here extends prior reduced metabolic systems by placing these metabolically relevant human tissue compartments within a shared circulating media environment and linking tissue-specific readouts to circuit-level functional and metabolomic endpoints.

A central advance of this study is the demonstration that shared perfusion acts as a state-setting variable for engineered human tissues. Under the Mid nutrient condition, multi-tissue interaction enriched tissue-aligned functional programs, whereas isolated tissues more often retained selected stress, inflammatory, injury, regenerative, or remodeling-associated programs. By preserving compartment-specific readouts within a shared circulating environment, the six-tissue format allows tissue-state remodeling and stress-propagation responses to be evaluated across connected human metabolic compartments. These findings support a broader principle for MPS design: cross-compartment signaling is an active culture variable that can shape engineered tissue phenotype ^36,37,87,88^.

The effect of tissue interaction depended on nutrient context. Under the Mid reference condition, shared perfusion supported compartment-aligned functional programs across the circuit. Under High nutrient exposure, the same tissue connectivity was associated with stronger inflammatory and nutrient-stress-associated gene expression in liver organoids and islets relative to matched isolated cultures. This distinction is important for interpreting the platform as a resource: tissue interaction shaped engineered tissue phenotype in a state-dependent, context-specific manner. The system therefore provides an experimental framework for studying conditions in which shared signaling supports functional specialization and conditions in which shared signaling amplifies stress-associated remodeling. This context dependence is relevant to metabolic disease biology, where endocrine and metabolite exchange contribute to homeostasis under physiological conditions and can propagate inflammatory, lipotoxic, proteostatic, or counter-regulatory signals during sustained metabolic stress ^11,46,86,89,90^.

The graded nutrient studies define a staged set of reference states for the interacting six-tissue circuit. Low conditions modeled selected features of lower nutrient availability and were associated with tissue-specific programs related to maintenance, substrate mobilization, epithelial surveillance, glucagon-responsive liver organoid signaling, and brain organoid proteostatic/redox regulation ^91–93^. Mid conditions modeled selected features of increased nutrient availability and induced anabolic, secretory, storage, bile-acid, gut nutrient-sensing, and brain nutrient-responsive programs, accompanied by increased insulin, C-peptide, albumin secretion, and remodeling of the shared extracellular metabolite environment ^45,61,86^. High nutrient exposure shifted the circuit toward stress-associated metabolic dysfunction, including persistent residual glucose accumulation, reduced net glucose depletion over time, elevated glucagon, proteostatic and lipid-redox stress programs, islet inflammatory and proteotoxic remodeling, intestinal stress activation, and brain organoid stress-associated transcriptional remodeling, consistent with high-fat nutrient stress ^10,80,94,95^.

The High nutrient state also demonstrates how the platform can connect tissue-specific stress programs to shared extracellular remodeling. *In vivo*, impaired hepatic lipid buffering can propagate lipid-derived stress signals and metabolic intermediates to distal tissues, contributing to systemic metabolic dysfunction ^89,90^. In the six-tissue MPS, High nutrient exposure coincided with liver organoid stress programs, altered bile-acid and lipid-associated metabolites in the shared perfusate, and stress-associated remodeling across distal tissue compartments. The enrichment of HIF-1α, inflammatory/stress, synaptic/neural, TGFβ/BMP, and matrix-associated programs in the brain organoid compartment is consistent with literature linking nutrient excess to altered brain mitochondrial, inflammatory, and synaptic pathways, as well as broader diabetes-associated *BMP*/matrix remodeling programs ^94,95^. Together, the Low, Mid, and High conditions provide reference perturbation states for comparing future tissue combinations, donor backgrounds, media formulations, metabolic stressors, or therapeutic interventions.

The drug studies demonstrate the utility of the multi-tissue resource for separating functional and transcriptional response modes under the same High nutrient stress state. Metformin most effectively preserved circuit-level net glucose depletion without a proportional increase in insulin or C-peptide accumulation, consistent with improved shared-media glucose handling without increased β-cell secretory burden over the measured window. Transcriptionally, metformin engaged liver organoid nutrient-response and reverse-cholesterol-transport programs, islet proteostatic and autophagy-associated remodeling, adipocyte lipid and energy-handling programs, and muscle-associated tissue-stabilizing pathways ^67,77,96^. Semaglutide produced broader remodeling in gut–brain–islet–liver compartments, including epithelial maintenance, bile-acid metabolism, glycogen-associated programs, endocrine stimulus-response pathways, and neurometabolic transcriptional programs, with more limited acute impact on net glucose depletion over the measured window. The platform therefore distinguished a metformin-associated functional glucose-handling response from a semaglutide-associated tissue-state remodeling response. These differences emphasize the value of multi-tissue MPS models for testing whether a therapy modifies a shared-media functional endpoint, reshapes cross-compartment signaling, or engages tissue-specific adaptive programs^34,60,72,97^.

Several measurement considerations define how this resource should be used. Net glucose depletion reflects disappearance of glucose from the shared perfusate and captures aggregate circuit-level glucose handling and not tissue-specific glucose uptake or metabolic flux. Insulin and C-peptide measurements were collected from shared circulating media after 48-hour intervals and therefore reflect accumulated endocrine output within a long-term interacting circuit rather than acute glucose-stimulated insulin secretion ^98^. Similarly, metabolomic readouts represent shared-media metabolite availability. These measurements are most useful for defining circuit-level extracellular state and nominating candidate metabolite axes for future isotope-tracing, compartment-specific uptake, or targeted perturbation studies.

This study has limitations typical of current multi-tissue MPS platforms. The model simplifies autonomic and neuroendocrine regulation, immune and vascular complexity, renal clearance, lymphatic transport, whole-body scaling, pharmacokinetics, gut peristalsis, contraction-dependent skeletal muscle metabolism, adipose tissue vascularization, and complete brain-mediated control of feeding and energy balance ^11,36,42^. These constraints are especially relevant for semaglutide, whose clinical effects depend on distributed gut–brain signaling, appetite regulation, gastric emptying, pharmacokinetics, and body-weight-associated mechanisms that are only partially represented in this *in vitro* circuit ^34,71,99^. Future iterations will extend the current platform from a state-resolving resource toward a mechanism-testing system.

Collectively, these data introduce AnthroHive and MOTIVE-6 as complementary human MPS technologies for studying selected features of glucose-regulatory tissue interaction under controlled shared perfusion. The six-tissue MPS revealed that multi-tissue context modifies engineered tissue state, graded nutrient exposure shifts the circuit from lower-nutrient maintenance toward compensatory remodeling and then stress-associated dysfunction, and metformin and semaglutide separate into distinct response modes under High nutrient conditions. By integrating functional readouts, endocrine signaling, tissue transcriptomics, and shared-media metabolomics across six metabolically relevant human tissue compartments, this platform provides a reusable framework for investigating how nutrient availability and therapeutic perturbation reshape circuit-level metabolic responses in human MPS models.

## STAR Methods

### Key resources table

| Reagent or Resource | Source | Identifier |
| --- | --- | --- |
| **Antibodies** |  |  |
| tyrosine hydroxylase | Santa Cruz Biotechnology | cat. no. sc25269 |
| insulin | Nordic | cat. no. GPASW-INS-FITC |
| zonula occludens-1 | Invitrogen | cat. no. 33-9111 |
| Mucin-2 | Santa Cruz Biotechnology | cat. no. sc-515032 AF594 |
| HCS LipidTox™ Neutral Lipid Stain | Invitrogen | cat. no. H34475 |
| Desmin D93F5 XP Rabbit | Cell Signaling | cat. no. 5332 |
| Alexa Fluor 594-conjugated secondary antibody, Donkey anti-mouse IgG | Thermo Fisher | cat. no. A-21203 |
| Alexa Fluor 488-conjugated secondary antibody, Donkey anti-Rabbit IgG | Abcam | cat. no. 150073 |
| **Biological Samples** |  |  |
| Human intestinal tissue | LifeNet Health | De-identified donor human tissue |
| Human liver tissue | LifeNet Health | De-identified donor human tissue |
| Human omental white pre-adipocytes | Lonza | cat. no. PT-5005 |
| Human skeletal muscle myoblasts | Lonza | cat. no. CC-2580 |
| Human pancreatic islets | Prodo Labs | De-identified donor islets |
| Human iPSC line - KOLF2.1J | Jackson Labs | cat. no. JIPSC001000 |
| L-WRN cells | ATCC | cat. no. CRL-3276 |
| **Chemicals, Peptides, and Recombinant Proteins** |  |  |
| ABS-M30 White material | Stratasys | cat. no. 333-60302 |
| QSR support material | Stratasys | cat. no. 333-63500 |
| Growth Factor Reduced Matrigel | Corning | cat. no. 356231 |
| hESC-Qualified Matrigel | Corning | cat. no. 354277 |
| Williams E Media | Gibco | cat. no. A12176-01 |
| DMEM (w/o glucose) | Gibco | cat. no. A14430-01 |
| DMEM/F-12 | Gibco | cat. no. 11320033 |
| Neurobasal medium | Gibco | cat. no. 21103049 |
| StemFlex™ Medium | Gibco | cat. no. A3349401 |
| Bovine Serum Albumin | Sigma-Aldrich | cat. no. A9418 |
| Donkey Serum | Sigma-Aldrich | cat. no. D9663 |
| Horse Serum | Gibco | cat. no. 16050-130 |
| KnockOut™ Serum Replacement | Gibco | cat. no. 10828028 |
| PGM-2 Preadipocyte Growth Medium-2 Bullet Kit | Lonza | cat. no. PT-8002 |
| SkGM-2 Skeletal Muscle Cell Growth Medium 2 Bullet Kit | Lonza | cat. no. CC-3245 |
| PIM(S) | Prodo Labs | cat. no. PIM-S001GMP |
| PIM(ABS) | Prodo Labs | cat. no. PIM-ABS001GMP |
| PIM(G) | Prodo Labs | cat. no. PIM-G001GMP |
| HCM Hepatocyte Culture Medium Bullet Kit | Lonza | cat. no. CC-3198 |
| HepatiCult™ Human Organoid Growth Medium | StemCell | cat. no. 100-0385 |
| HepatiCult™ Human Organoid Differentiation Medium | StemCell | cat. no. 100-0383 |
| Dexamethasone | ChemCruz | cat. no. sc-29059 |
| Dorsomorphin | Sigma Aldrich | cat. no. P5499 |
| Metformin | Tokyo Chemistry Industry | cat. no. M2009 |
| Semaglutide | AdipoGen | cat. no. AG-CP3-0040 |
| IWP-2 | BioGems | cat. no. 6866167 |
| cAMP | BioGems | cat. no. 6099240 |
| Laminin | Corning | cat. no. 354232 |
| Heparin | Sigma Aldrich | cat. no. H3149-25KU |
| D-(+)-Glucose | Sigma-Aldrich | cat. no. G7021 |
| Lipid Mix | Sigma-Aldrich | cat. no. L0288 |
| GlutaMAX | Gibco | cat. no. 35050061 |
| HEPES | Gibco | cat. no. 15630-080 |
| Non-Essential Amino Acids | Gibco | cat. no. 11140-050 |
| Penicillin /Streptomycin | Gibco | cat. no. 15140122 |
| Nicotinimide | Sigma-Aldrich | cat. no. N0636 |
| Ascorbic acid | Sigma Aldrich | cat. no. A4544 |
| N-Acetylcysteine | Millipore Sigma | cat. no. 45-A7250 |
| SAG | BioGems | cat. no. 9128694 |
| A83-01 | BioGems | cat. no. 9094360 |
| Y27632 | BioGems | cat. no. 1293823 |
| SB202190 | BioGems | cat. no. 1523072 |
| B-27 | Gibco | cat. no. 17504-001 |
| B-27 without Vitamin A | Gibco | cat. no. 12587010 |
| N-2 | Gibco | cat. no. 17502048 |
| 2-Mercaptoethanol | Gibco | cat. no. 21985023 |
| CHIR99021 | StemCell | cat. no. 72054 |
| Gastrin | Anaspec- Eurogentec | cat. no. AS-64149 |
| Prostaglandin E2 | BioGems | cat. no. 3632464 |
| Recombinant Human Insulin | Sigma Aldrich | cat. no. 91077C |
| Recombinant Human HGF | PeproTech | cat. no. 100-39 |
| Recombinant Human EGF | Gibco | cat. no. PHG0313 |
| Recombinant Human bFGF | PeproTech | cat. no. 100-18B |
| Recombinant Human FGF-8 | PeproTech | cat. no. 100-25 |
| Recombinant Human BDNF | PeproTech | cat. no. 450-02 |
| Recombinant Human GDNF | PeproTech | cat. no. 450-10 |
| Recombinant Human Noggin | PeproTech | cat. no. 120-10C |
| NucBlue | Invitrogen | cat. no. R37605 |
| ActinRed | Invitrogen | cat. no. R37112 |
| Vectashield HardSet Antifade Mounting Medium | Vector Laboratories | cat. no. H-1500-10 |
| Cell Recovery Solution | Corning | cat. no. 354270 |
| RNeasy Mini kit | Qiagen | cat. no. 74106 |
| Critical Commercial Assays |  |  |
| Albumin ELISA | RayBio | cat. no. ELH-Albumin |
| Glucose ELISA | Invitrogen | cat. no. EIAGLUC |
| Human Metabolic Luminex Performance Premixed Kit | R&D Systems | cat. no. FCSTM19-09 |
| Human Luminex Discovery Assay | R&D Systems | cat. no. LXSAHM-05 |
| Milliplex Human Neuroscience Magnetic Bead Panel 1 | Millipore | cat. no. HNS1MAG-95K |
| Deposited Data |  |  |
| Raw RNAseq Data | GEO: | Accession no. GSE330098 |
| Raw Metabolomics Data | MetaboLights: | Accession no. MTBLS14530 |
| Software and Algorithms |  |  |
| Luminex xPONENT for Luminex 200 v4.2 | Luminex Corporation | https://us.diasorin.com/en/luminex |
| ProcartaPlex Analysis App | Thermo Fisher Scientific | https://apps.thermofisher.com/apps/procartaplex |
| NIS-Elements | Nikon Instruments | https://www.microscope.healthcare.nikon.com/products/software/nis-elements |
| FIJI/Image J | National Institutes of Health | https://imagej.net/software/fiji |
| Nextflow, v24.04.4 | Nextflow contributors / Seqera Labs | Di Tommaso et al., 2017; https://www.nextflow.io |
| SortMeRNA, v4.3.7 | SortMeRNA developers | Kopylova et al., 2012; https://github.com/sortmerna/sortmerna |
| STAR, v2.7.11b | STAR developers | Dobin et al., 2013; https://github.com/alexdobin/STAR |
| Salmon, v1.10.3 | COMBINE-lab | Patro et al., 2017; https://combine-lab.github.io/salmon/ |
| R, v4.4.2 | R Foundation for Statistical Computing | R Core Team, 2024; https://www.R-project.org/ |
| Rstudio IDE | Posit Software, PBC | Posit team, 2025; https://posit.co/products/open-source/rstudio |
| DESeq2, v1.46 | Bioconductor | Love et al., 2014 |
| sva, v3.54.0 | Bioconductor | Leek et al., 2012 |
| ComBat | Bioconductor | Johnson et al., 2007; Leek et al., 2012 |
| limma v3.62.2 | Bioconductor | Ritchie et al., 2015 |
| fgsea, v1.32 | Bioconductor | Korotkevich et al., 2019; https://bioconductor.org/packages/fgsea |
| GSEA, v 2.2.3. | Broad Institute | http://software.broadinstitute.org/gsea/ |
| MSigDB | Broad Institute / UC San Diego | Liberzon et al., 2011; Subramanian et al., 2005; https://www.gsea-msigdb.org/gsea/msigdb |
| Cytoscape v3.7.1 | Cytoscape Consortium | Shannon et al., 2003; https://cytoscape.org/ |
| EnrichmentMap | EnrichmentMap developers | Merico et al., 2010; https://enrichmentmap.org/ |
| AutoAnnotate | AutoAnnotate developers | Kucera, 2016 |
| BulkSignalR, v1.0.5 | BulkSignalR developers | Villemin et al., 2023; https://github.com/jcolinge/BulkSignalR |
| Morpheus | Broad Institute | https://software.broadinstitute.org/morpheus/ |
| MZmine, v3.53 | MZmine development team | Schmid et al., 2023; https://mzmine.org/ |
| MetaboAnalyst 6.0 | McGill University / Xia Lab | Pang et al., 2024; Xia et al., 2010; https://www.metaboanalyst.ca/ |
| mixOmics v6.30 | mixOmics developers | Rohart et al., 2017; https://mixomics.org/ |
| GraphPad Prism, v11.0 | GraphPad Software | https://www.graphpad.com/scientificsoftware/prism/ |
| BioRender | BioRender | https://www.biorender.com/ |

### Lead Contact and Materials Availability

Information and requests for resources and reagents should be directed to and will be fulfilled by the lead contact, Martin Trapecar.

### Experimental model and study participant details

<u>Human pancreatic islets</u> - Human islets from two deidentified non-diabetic donors were obtained from Prodo Laboratories (Aliso Viejo, CA, USA). Upon receipt, islets were recovered in PIM(S) supplemented with PIM(ABS) and PIM(G) (Prodo Laboratories; cat. no. PIM-S001GMP, cat. no. PIM-ABS001GMP, and cat. no. PIM-G001GMP) and distributed into permeable cell culture inserts (CELLTREAT, 230635) at approximately 100 islet equivalents (IEQ) per insert.

<u>Human adipocyte cultures</u> - Human visceral pre-adipocytes (Lonza, cat. no. PT-5005) were cultured according to the manufacturer’s instructions using PGM-2 Preadipocyte Growth Medium-2 Bullet Kit (Lonza, cat. no. PT-8002). Cells were seeded onto permeable cell culture inserts (CELLTREAT, cat. no. 230635) coated with 2% Matrigel at approximately 1 × 10^5 cells per insert and differentiated into mature adipocytes for 10 days before the start of experiments.

<u>Human skeletal muscle cultures</u>- Human skeletal muscle myoblasts (Lonza, cat. no. CC-2580) were cultured according to the manufacturer’s instructions using SkGM-2 Skeletal Muscle Cell Growth Medium-2 BulletKit (Lonza, cat. no. CC-3245). Cells were seeded onto permeable cell culture inserts (CELLTREAT, cat. no. 230635) coated with 2% Matrigel at approximately 1 × 10^5 cells per insert and differentiated into multinucleated myotubes with 2 % horse serum (Gibco, cat. no. 16050-130) for 10 days before the start of experiments.

<u>Human gut epithelial cultures</u>- Human colon tissue was obtained from LifeNet Health LifeSciences and processed immediately upon arrival. Large-intestinal epithelial stem cells were isolated and expanded as colon organoids in WRN-based medium as previously described ^100,101^. For WRN-conditioned medium production, L-WRN fibroblast-like cells (ATCC, cat. no. CRL-3276) were expanded in DMEM-based growth medium grown until over-confluent, rinsed, and cultured in fresh medium for serial 24-hour collections of Wnt-3A-, R-spondin-, and Noggin-enriched conditioned medium.

Tissue was digested in 2 mg ml−1 collagenase I (StemCell, cat. no. 07416) for 40 min at 37°C followed by mechanical dissociation, and isolated crypts were resuspended in growth factor-reduced Matrigel (Corning, cat. no. 356231) and polymerized at 37°C. Organoids were grown in expansion medium consisting of Advanced DMEM/F12 supplemented with L-WRN conditioned medium (65% vol/vol, ATCC, cat. no. CRL-3276), 2 mM Glutamax (Gibco, cat. no. 35050061), 10 mM HEPES (Thermo Fisher, cat. no. 15630–080), Penicillin/Streptomycin (Gibco, cat. no. 15140122), 50 ng/mL EGF (Gibco, cat. no. PHG0313), N2 supplement (Gibco, cat. no. 17502048), B-27 Supplement (Gibco, cat. no. 17504-001), 1 nM human gastrin (Anaspec- Eurogentec, cat. no. AS-64149), 0.5 mM N-acetyl cysteine (Sigma, cat. no. 45-A7250), 10 mM nicotinamide (Sigma, cat. no. N0636) 10 µM SB202190 (BioGems, cat. no. 1523072), 500 nM A83–01 (BioGems, cat. no. 9094360), 5 nM prostaglandin E2 (BioGems, cat. no. 3632464) at 37°C and 5% CO2. Organoids were passaged every 7 days by incubating in Cell Recovery Solution (Corning, cat. no. 354253) for 40 min at 4 °C, followed by mechanical dissociation and reconstitution in fresh Matrigel at a 1:4 ratio.

For experiments, gut organoids were removed from Matrigel, dissociated with TrypLE into a single-cell suspension, and seeded onto 2% Matrigel-coated permeable inserts (CELLTREAT, cat. no. 230635) at approximately 1 × 10^5 cells per insert to generate epithelial monolayers. Gut barrier maintenance was assessed by transepithelial electrical resistance (TEER) every other day.

<u>Human liver organoid cultures</u> - Human liver tissue was obtained from LifeNet Health LifeSciences and processed immediately upon arrival. Hepatic cells were isolated and expanded as liver organoids as previously described ^102^. Tissue was mechanically minced and enzymatically digested to isolate hepatic cells, which were resuspended in growth factor-reduced Matrigel (Corning, cat. no. 356231) and polymerized at 37°C.

Organoids were maintained in HepatiCult Human Organoid Growth Medium (STEMCELL Technologies, cat. no. 100-0385) for expansion. Organoids were passaged every 7 days by incubating in Cell Recovery Solution (Corning, cat. no. 354253) for 40 min at 4 °C, followed by mechanical dissociation and reconstitution in fresh Matrigel at a 1:4 ratio and seeded onto permeable transwell inserts for experimental use. Prior to experiments, organoids were differentiated for 10 days in HepatiCult Human Organoid Differentiation Medium (STEMCELL Technologies, cat. no. 100-0383) supplemented with dexamethasone (ChemCruz, cat. no. sc-29059). Hepatocyte differentiation and functional maturation was evaluated by albumin secretion.

<u>Human midbrain organoids</u> - Midbrain organoids were generated from the KOLF2.1J hiPSC line (The Jackson Laboratory, cat. no. JIPSC001000) as previously described ^103,104^. Briefly, approximately 10,000 iPSCs were dissociated and aggregated as embryoid bodies in ultra-low-attachment U-bottom 96-well plates in Embryoid Body Medium. On days 1 and 3, medium was half changed with Midbrain Induction Medium (MIM) containing 2 μM dorsomorphin, 2 μM A83-01, 1 μM IWP2, and 3 μM CHIR99021. Patterning was continued on day 4 with MIM supplemented with 100 ng/mL FGF8 (PeproTech, cat. no. 100-25) and 2 μM SAG (BioGems, cat. no. 9128694), and on day 7 with MIM supplemented with 100 ng/mL FGF8, 2 μM SAG, 2.5 μg/mL insulin (Sigma Aldrich, cat. no. 91077C), and 200 ng/mL laminin (Corning, cat. no. 354232). Embryoid bodies were embedded on day 7 in 30 μL growth factor-reduced Matrigel droplets, transferred to suspension culture, and maintained from day 9 onward in Midbrain Maturation Medium containing 10 ng/mL BDNF (Peprotech, cat. no. 450-02), 10 ng/mL GDNF (Peprotech, cat. no. 450-10), 200 μM ascorbic acid (Sigma Aldrich, cat. no. A4544), and 125 μM cAMP (BioGems, cat. no. 6099240). Organoids were matured for approximately 3 months before use in multiorgan studies. For MPS experiments, individual midbrain organoids were transferred to permeable 24-well inserts (CELLTREAT, cat. no. 230635), overlaid with 100 μL growth factor-reduced Matrigel, incubated at 37°C for 30 min, and maintained in maturation medium until integration into the system.

### Method details

#### Device design, fabrication and preparation

To support controlled recirculating perfused culture for six-tissue interaction studies, we developed the AnthroHive platform together with Multiorgan Tissue Interaction Vessels-6 (MOTIVE-6). The AnthroHive housing was 3D printed at the Johns Hopkins All Children’s Hospital Center for Medical Simulation and Innovative Education from custom design files. The housing was fabricated using a Stratasys F370 printer with ABS White material (Stratasys, cat. no. 333-60302) and QSR support material (Stratasys, cat. no. 333-63500). During post-processing, support material was removed, and the printed housing was cleaned in an Omegasonics 1900BT Ultrasonic Cleaner for 20 min using Stratasys WaterWorks P400SC solution. The housing was then rinsed under flowing water for 15 min and allowed to air dry. The AnthroHive housing does not directly contact culture media and serves as an incubator-compatible support platform for pump-mounted MOTIVE-6 operation.

The AnthroHive platform is configured to house nine peristaltic pumps (Takasago RK-QII1.5S) operated by a custom pump controller manufactured by RSEN s.p. (Slovenia) for controlled media recirculation through MOTIVE-6 vessels. For the experiments described here, pumps were operated within a flow-rate range of 0.5–3 mL/min. Each MOTIVE-6 vessel contains six interconnected compartments designed to accommodate organoids or permeable membrane inserts. Each compartment was designed to accept a 24-well-format 0.4 µm pore-size permeable membrane insert, allowing tissue modules to remain physically separated while sharing a common recirculating medium. Each assembled MOTIVE-6 vessel contained 8 mL of shared circulating medium. MOTIVE-6 vessel bodies were CNC machined from natural polyether ether ketone (PEEK) with an as-machined finish by Fictiv (San Francisco, CA, USA). MOTIVE-6 lids were 3D printed at the Johns Hopkins All Children’s Hospital Simulation Center using ABS White material on a Stratasys F370 printer.

Before experimental use, MOTIVE-6 vessels, lids, and connectors were sterilized using a V10 Air Plasma Sterilizer (Plasma Bionics) and handled under sterile conditions during MPS assembly. Sterilized MOTIVE-6 vessels were fluidically connected using polypropylene luer-lock connectors to establish sterile closed-loop media recirculation between vessel compartments and the AnthroHive pump system. For all six-tissue experiments, compartment positions within MOTIVE-6 were kept constant across vessels to preserve vessel layout, flow path, and tissue placement.

#### Graded nutrient medium formulation

A shared experimental medium was used for multiorgan interaction studies. The medium base consisted of a 1:1 mixture of Williams E Medium (Gibco, cat. no. A12176-01) and glucose-free DMEM (Gibco, cat. no. A14430-01), providing a basal glucose concentration of 5.55 mM. The shared medium was further supplemented with 1× GlutaMAX (Gibco, cat. no. 35050061), 10 mM HEPES (Gibco, cat. no. 15630-080), 0.1% bovine serum albumin (Sigma-Aldrich, cat. no. A9418), 1 mM N-acetylcysteine (MilliporeSigma, cat. no. A7250), 2 μM CHIR99021 (STEMCELL Technologies, cat. no. 72054), 20 ng/mL HGF (PeproTech, cat. no. 100-39), 50 ng/mL Noggin (PeproTech, cat. no. 120-10C), and GA-1000, rhEGF, ascorbic acid, and transferrin (Lonza, cat. no. CC-4182), each added at 0.1% of the supplied stock. Full media formulation is described in Table S1.

Nutrient conditions were generated by supplementing shared culture medium with D-(+)-glucose (Sigma-Aldrich, cat. no. G7021) and Lipid Mixture 1 (Sigma-Aldrich, L0288). Lipid Mixture 1 is a complex, non-animal-derived lipid formulation containing arachidonic acid (2 μg/mL), linoleic acid, linolenic acid, myristic acid, oleic acid, palmitic acid, and stearic acid (10 μg/mL each), cholesterol from New Zealand sheep’s wool (0.22 mg/mL), Tween-80 (2.2 mg/mL), tocopherol acetate (70 μg/mL), and Pluronic F-68 (100 mg/mL), solubilized in cell-culture-grade water with 10% ethanol by volume. Low, Mid, and High nutrient media contained 6, 12, or 18 mM glucose with 0.5%, 2%, or 4% Lipid Mixture 1, respectively, expressed as v/v dilution of the supplied stock, shown in Table S2)

#### MOTIVE-6 interaction experiments

MOTIVE-6 vessels were equilibrated in complete Mid medium for 24 h before the start of each experiment. In parallel, tissues maintained in transwells were adapted for 24 h in a 1:1 mixture of Mid medium and their respective tissue-specific maintenance medium, followed by an additional 24 h equilibration in 100% Mid medium. This staged equilibration was used to reduce carryover of residual glucose, insulin, and tissue-specific supplements while minimizing acute perturbation during transition into the shared experimental medium.

At the start of each experiment, vessels and tissues were washed twice with DPBS. Each MOTIVE-6 vessel received 8 mL of the appropriate experimental nutrient medium representing Low (6 mM glucose + 0.5% lipid), Mid (12 mM glucose + 2% lipid), or High (18 mM glucose + 4% lipid) nutrient conditions. For drug-treatment studies performed under High nutrient conditions, media were supplemented with semaglutide (130 ng/mL, AdipoGen, cat. no. AG-CP3-0040) or metformin (1 μg/mL, TCI, cat. no. M2009).

Equilibrated tissues were assembled into MOTIVE-6 in the order shown in Fig.1B, and tissue positions were kept consistent across experiments. Vessels were connected to AnthroHive pump heads, with flow rate of 3 mL/min and maintained at 37°C, 5% CO2, and 95% humidity throughout the experiment.

Isolation controls consisted of the same tissue types cultured separately in transwells or plates under matched nutrient conditions. Each MOTIVE-6 vessel or matched isolation culture constituted one biological replicate. Experiments were performed across two independent runs, yielding n = 6 for Mid, n = 4 each for Low, High, metformin, and semaglutide conditions, and n = 3 for isolation controls. Circulating media were collected and replenished in full every 2 days over a total 4-day experiment. At days 0, 2, and 4, tissues were imaged by brightfield microscopy, and gut barrier integrity was measured by TEER. On day 4, tissues were harvested for downstream analyses.

Matrigel-embedded tissues were released using Cell Recovery Solution (Corning, cat. no. 354270), washed with DPBS, lysed in RLT buffer (Qiagen, cat. no. 74106) and stored at −80 °C for RNA extraction. All other tissues were washed with DPBS and directly lysed in RLT buffer before storage at −80 °C.

#### Immunostaining and imaging

For immunofluorescence staining, samples were washed with DPBS and fixed in 4% paraformaldehyde for approximately 15 min. Samples were washed with DPBS, permeabilized with 0.1% Triton X-100 for 10 min, and blocked in DPBS containing 2% bovine serum albumin for 2 h. Primary antibodies were diluted in DPBS containing 1% bovine serum albumin and 0.1% Tween-20 and incubated for 1–2 h at room temperature. Samples were then washed with DPBS containing 0.1% Tween-20 and incubated with fluorophore-conjugated secondary antibodies, where required. Nuclear, actin, and neutral lipid stains were used as indicated.

Primary antibodies and directly conjugated staining reagents included anti-tyrosine hydroxylase antibody (Santa Cruz Biotechnology, cat. no. sc-25269), FITC-conjugated anti-insulin antibody (Nordic, cat. no. GPASW-INS-FITC), FITC-conjugated anti-zonula occludens-1 antibody (Thermo Fisher Scientific, cat. no. 33-9111), Alexa Fluor 594-conjugated anti-Mucin-2 antibody (Santa Cruz Biotechnology, cat. no. sc-515032 AF594), anti-desmin D93F5 XP rabbit monoclonal antibody (Cell Signaling Technology, cat. no. 5332), and HCS LipidTOX Neutral Lipid Stain (Thermo Fisher Scientific, cat. no. H34475). Alexa Fluor 594-conjugated donkey anti-mouse IgG secondary antibody (Thermo Fisher Scientific, cat. no. A-21203) was used for anti-tyrosine hydroxylase. Alexa Fluor 488-conjugated donkey anti-rabbit IgG secondary antibody (Abcam, cat. no. ab150073) was used for anti-desmin staining.

For muscle samples, blocking was performed in DPBS containing 1% donkey serum (Sigma-Aldrich, cat. no. D9663) and 0.1% Tween-20 overnight at 4°C, followed by incubation with anti-desmin primary antibody overnight at 4°C. Nuclear staining was performed using NucBlue ReadyProbes Reagent (Thermo Fisher Scientific, cat. no. R37605), actin staining with ActinRed 555 ReadyProbes Reagent (Thermo Fisher Scientific, cat. no. R37112), and neutral lipid staining with HCS LipidTOX Neutral Lipid Stain.

Samples were mounted using VECTASHIELD HardSet Antifade Mounting Medium and imaged on a Nikon Eclipse Ti2 microscope equipped with an A1R confocal scan head using NIS-Elements software. Images were acquired using a 20× dry objective (Plan Apo 20×/0.75, OFN25) or a 40× oil-immersion objective (Plan Fluor 40×/1.30 Oil, cover glass correction 0.17 mm). Fluorophores were excited using 405 nm, 488 nm argon, 561 nm DPSS, and/or 633 nm HeNe laser lines, and images were collected as single optical sections or z-stacks, as appropriate. For comparisons across experimental groups, acquisition settings were kept constant within each staining panel, and representative images were processed in FIJI ^105^ uniformly for brightness and contrast.

#### Conditioned media biomarker measurements

**Luminex assays** - Circulating conditioned media were analyzed for metabolic, inflammatory, and endocrine biomarkers using multiplex bead-based Luminex assays according to the manufacturers’ instructions. A human metabolic panel (R&D Systems, cat. no. FCSTM19-09) was used to quantify C-peptide, MCP-1, GLP-1, glucagon, IL-6, insulin, leptin, peptide YY, and TNF-α. A human discovery panel (R&D Systems, cat. no. LXSAHM-05) was used to measure adiponectin, C-reactive protein, IL-8, resistin, and PAI-1. To assess biomarkers associated with neurological dysfunction in the brain organoid apical compartment, a Milliplex Human Neuroscience Magnetic Bead Panel (Millipore, cat. no. HNS1MAG-95K) was used to quantify GFAP, UCHL1, DJ1, NSE, α-synuclein, and TGM2 according to the manufacturer’s instructions. Plates were read on a Luminex 200 instrument. Median fluorescent intensity values were background corrected and averaged across technical replicates, and analyte concentrations were calculated in xPONENT software using 5-parameter logistic standard curves.

#### Glucose colorimetric assay

Glucose concentration in conditioned media was measured using a colorimetric assay kit (Invitrogen, cat. no. EIAGLUC) according to the manufacturer’s instructions. Absorbance was measured at 560 nm on a BioTek Cytation 1 plate reader. Percent glucose consumption was calculated relative to unconditioned base medium.

#### Human albumin ELISA

Albumin secretion was quantified in conditioned media using a human albumin ELISA kit (RayBiotech, cat. no. ELH-Albumin) according to the manufacturer’s instructions. Absorbance was measured at 450 nm on a BioTek Cytation 1 plate reader.

### RNA analysis

#### RNA purification and sequencing

RNA was isolated from harvested tissues using the RNeasy Mini Kit (Qiagen, cat. no. 74106). Tissue lysates were prepared in RLT buffer supplemented with β-mercaptoethanol (Gibco, cat. no. 21985023) and stored at −80°C until purification. RNA concentration and purity were assessed using NanoDrop spectrophotometry and Qubit fluorometry, and RNA integrity was evaluated using Agilent RNA ScreenTape on a 4200 TapeStation. Bulk RNA-seq libraries were prepared by Novogene (Sacramento, CA) using poly(A)+ mRNA enrichment and standard Illumina library construction protocols. Libraries were sequenced on an Illumina platform to generate paired-end reads.

#### RNA-seq read processing and differential expression analysis

Raw FASTQ files were processed using the nf-core/rnaseq pipeline (v3.19) executed in Nextflow, v24.04.4 ^106^. Reads were aligned to the human reference genome GRCh38.p14, NCBI RefSeq assembly GCF_000001405.40, using STAR, v2.7.11b ^107^. Residual ribosomal RNA reads were identified using SortMeRNA, v4.3.7 ^108^, and transcript quantification was performed with Salmon, v1.10.3 ^109^. Read processing was conducted on the DISCOVERY High Performance Computing Facility at Johns Hopkins University.

Transcript-level abundance estimates generated by Salmon were imported into R, v4.4.2 ^110^ using tximport and summarized to gene-level counts. Genes with zero counts in more than 50% of samples were removed prior to downstream analysis. Differential expression analysis was performed using DESeq2, v1.46 ^111^. Surrogate variable analysis was performed using svaseq from sva, v3.54.0 ^112^ as an exploratory assessment of latent technical variation. In this analysis, nutrient condition was defined as the variable of interest, and svaseq was used to estimate the number of surrogate variables explaining residual variation not captured by nutrient condition. This assessment indicated that independent experimental run was a major source of technical variation; therefore, differential expression models included experimental run as a batch covariate. Pairwise comparisons were carried out across nutrient, drug-treatment, and culture-context comparisons within each tissue type. P values were adjusted for multiple testing using the Benjamini-Hochberg method. For visualization only, batch effects were reduced using removeBatchEffect from limma v3.62.2 ^113^.

#### Transcriptomic pathway enrichment, clustering, and ligand-receptor analysis

For transcriptomic pathway analysis, fgsea, v1.32 92, was applied to preranked gene lists generated from DESeq2 results. Genes were ranked using −log10(P value) × log2(fold change), such that ranking incorporated both effect size and statistical evidence ^114–116^. Gene Ontology Biological Process (GO-BP) gene sets were obtained from MSigDB, version 2026.1.Hs 93,94. Pathways with nominal P < 0.05 in at least one tissue or contrast were retained for downstream visualization ^117–119^.

To compare pathway behavior across tissues and contrasts, normalized enrichment scores were converted into ΔNES values and assembled into a pathway-by-condition matrix. Missing values were set to 0, and the matrix was hierarchically clustered in Morpheus using one minus Pearson correlation. Meta-program clusters were defined from third-level dendrogram branches and semantically annotated.

Ligand-receptor interactions were inferred using BulkSignalR, v1.0.5 ^120^. Predicted interactions were summarized at the pathway level using the Reactome-derived database implemented in BulkSignalR. Statistical significance was assessed using q values, and interaction profiles were compared between tissues cultured in isolation and in multiorgan interaction to identify context-dependent crosstalk patterns. To identify interaction-associated ligand-receptor signatures, ligand-receptor pairs were further filtered for interactions detected in multiorgan culture but absent in isolation and grouped into functional signaling categories based on ligand-receptor annotations.

### Metabolomics analysis

#### Metabolomics sample preparation and UHPLC-HRMS acquisition

For metabolomics analyses, all sample preparation steps were carried out on ice. Briefly, 20 μL of conditioned media was extracted with 80 μL of pre-cooled methanol extraction solvent that had been chilled at −80°C for at least 1 h before use. Following solvent addition, samples were vortexed and centrifuged at 18,800 × g for 10 min at 0°C, then incubated at −80°C for 30 min to improve metabolite extraction. Samples were subsequently centrifuged again at 18,800 × g for 10 min at 4°C, and the supernatants were collected, dried, and reconstituted in 20 μL of 80% methanol containing 10% of a stable isotope-labeled metabolomics quality control mixture.

Untargeted UHPLC-HRMS metabolomics was performed using a Vanquish UHPLC coupled to a Q Exactive HF quadrupole-Orbitrap mass spectrometer (Thermo Scientific). Chromatographic separation was achieved using an Atlantis Premier BEH Z-HILIC VanGuard FIT column (2.1 mm × 150 mm, 2.5 μm particle size; Waters). Mobile phase A consisted of 10 mM ammonium carbonate with 0.05% ammonium hydroxide in water, and mobile phase B consisted of 100% acetonitrile. The gradient was as follows: 80% B at the start, a linear gradient from 80% to 20% B over 13 min, a 2 min hold at 20% B, return to 80% B in 0.1 min, and re-equilibration for 4.9 min, for a total run time of 20 min. The flow rate was 0.250 mL/min, the autosampler was maintained at 5°C, the column was maintained at 30°C, and the injection volume was 2 μL. Samples were analyzed separately in positive and negative electrospray ionization modes over an m/z range of 65–900. Pooled samples were subjected to data-dependent MS/MS acquisition to support metabolite confirmation.

#### Metabolomics data processing and statistical analysis

Raw metabolomics data were processed in MZmine, v3.53 using a batch workflow that included centroid mass detection, ADAP chromatogram building, smoothing, deconvolution by local minimum search, isotopic peak grouping, peak alignment, gap filling, duplicate peak filtering, blank feature filtering, custom database searching, adduct and complex searching, row filtering, and peak list export ^121^. Key processing parameters included a minimum chromatographic group size of 5 scans, minimum highest intensity of 1.0E4, deconvolution with a 95% chromatographic threshold, 0.05 min minimum search range in retention time, 10% minimum relative height, 1.0E4 minimum absolute height, peak duration of 0.05–5 min, isotopic grouping with 10 ppm m/z tolerance and 0.25 min retention time tolerance, and custom database matching against an in-house library with 10 ppm m/z tolerance and 0.3 min retention time tolerance.

Positive and negative ionization mode datasets were processed separately and subsequently merged in R, v4.4.2 ^110^. Peaks/features detected in fewer than half of samples were removed, and features without metabolite annotation were excluded from downstream statistical analysis. Metabolite annotations were assigned by matching accurate mass and retention time to an in-house metabolite library. Redundant metabolite identifications were resolved by averaging peak intensities across duplicate features. Peak intensities were median-normalized across samples, and experimental batch effects were corrected using ComBat from the sva package ^112,122^. Normalized and batch-corrected intensities were log2-transformed before statistical analysis and visualization. Because metabolites were measured from shared circulating media, comparisons were performed at the whole-system level.

Differential metabolite abundance was assessed in R using t-tests on log2-transformed, normalized, batch-corrected intensities, followed by Benjamini-Hochberg correction for multiple testing. Metabolomics comparisons were performed across nutrient and drug-treatment conditions at the shared-media, whole-system level. Tissue-specific comparisons were not performed because metabolites were measured from the shared circulating media. Partial least squares discriminant analysis (PLS-DA) was performed using mixOmics v6.30 to identify metabolites driving separation between experimental groups ^123^. Metabolite importance for the first two discriminant functions was assessed based on the magnitude of discriminant function loadings. To identify disease-associated metabolic signatures, significantly altered metabolites were analyzed in MetaboAnalyst 6.0 using metabolite set enrichment analysis with over-representation analysis against curated disease-associated metabolite set libraries ^124,125^.

### Quantification and statistical analysis

#### Replicate definition and experimental units

For multiorgan interaction experiments, one fully assembled MOTIVE-6 vessel containing all six tissue compartments was treated as one biological replicate. Interaction experiments included n = 4 biological replicate vessels for Low nutrient, High nutrient, High + metformin, and High + semaglutide conditions, and n = 6 biological replicate vessels for the Mid nutrient condition. For isolation experiments, one independently cultured tissue preparation maintained under the corresponding nutrient condition was treated as one biological replicate. Experiments were performed across two independent experimental runs. Technical replicates from plate-based assays, repeated assay wells, or imaging fields were averaged within each biological replicate before statistical analysis. For longitudinal media measurements, repeated measurements collected from the same MOTIVE-6 vessel over time were treated as repeated measures. For transcriptomic analyses, each harvested tissue compartment from an individual MOTIVE-6 vessel or matched isolation culture represented one RNA-seq sample for that tissue and condition. For metabolomics analyses, each circulating media sample represented the shared extracellular environment of one MOTIVE-6 vessel and was therefore analyzed at the whole-system level.

#### General statistical reporting

Statistical analyses for ELISA, Luminex, and TEER datasets were performed using GraphPad Prism v11.0 (GraphPad Software). Depending on the experimental design, statistical comparisons were performed where appropriate using unpaired t-test, one-way ANOVA, or two-way ANOVA with repeated measures, followed by Tukey’s multiple-comparisons test. Unless otherwise stated, multiple testing correction for omics analyses was performed using the Benjamini-Hochberg method. Statistical details for each experiment, including the number of biological replicates, statistical tests, and significance thresholds, are provided in the corresponding figure legends where applicable.

## Data and code availability

Bulk RNA-seq and metabolomics data generated in this study have been deposited in publicly available repositories. Gene expression matrices for our bulk RNA-seq assay are available through the Gene Expression Omnibus under accession number GSE330098. Raw RNAseq sequences can be found at the Sequence Read Archive (SRA, BioProject PRJNA1462736). Metabolomics mzXML spectra files are available through MetaboLights under accession number MTBLS14530. Code used for multi-omics analysis is available at https://github.com/oospina/Tissue_On_Chip_Multi_Omics_Analysis. Any additional information required to reanalyze the data reported in this paper is available from the lead contact upon reasonable request.

## Acknowledgments

This work has been supported in part by the Proteomics and Metabolomics Core Facility at the H. Lee Moffitt Cancer Center & Research Institute, an NCI designated Comprehensive Cancer Center (P30-CA076292). The study was supported in part by grants from NIGMS 5R35GM146900.

## Declaration of generative AI and AI-assisted technologies in the writing process

During the preparation of this work the author(s) used ChatGPT 5.5 (OpenAI) and Claude 4.5 (Anthropic) in order to support language refinement, grammar correction, and editorial clarity. After using this tool/service, the author(s) reviewed and edited the content as needed and take(s) full responsibility for the content of the published article.

---

**Supplementary Table S1:**
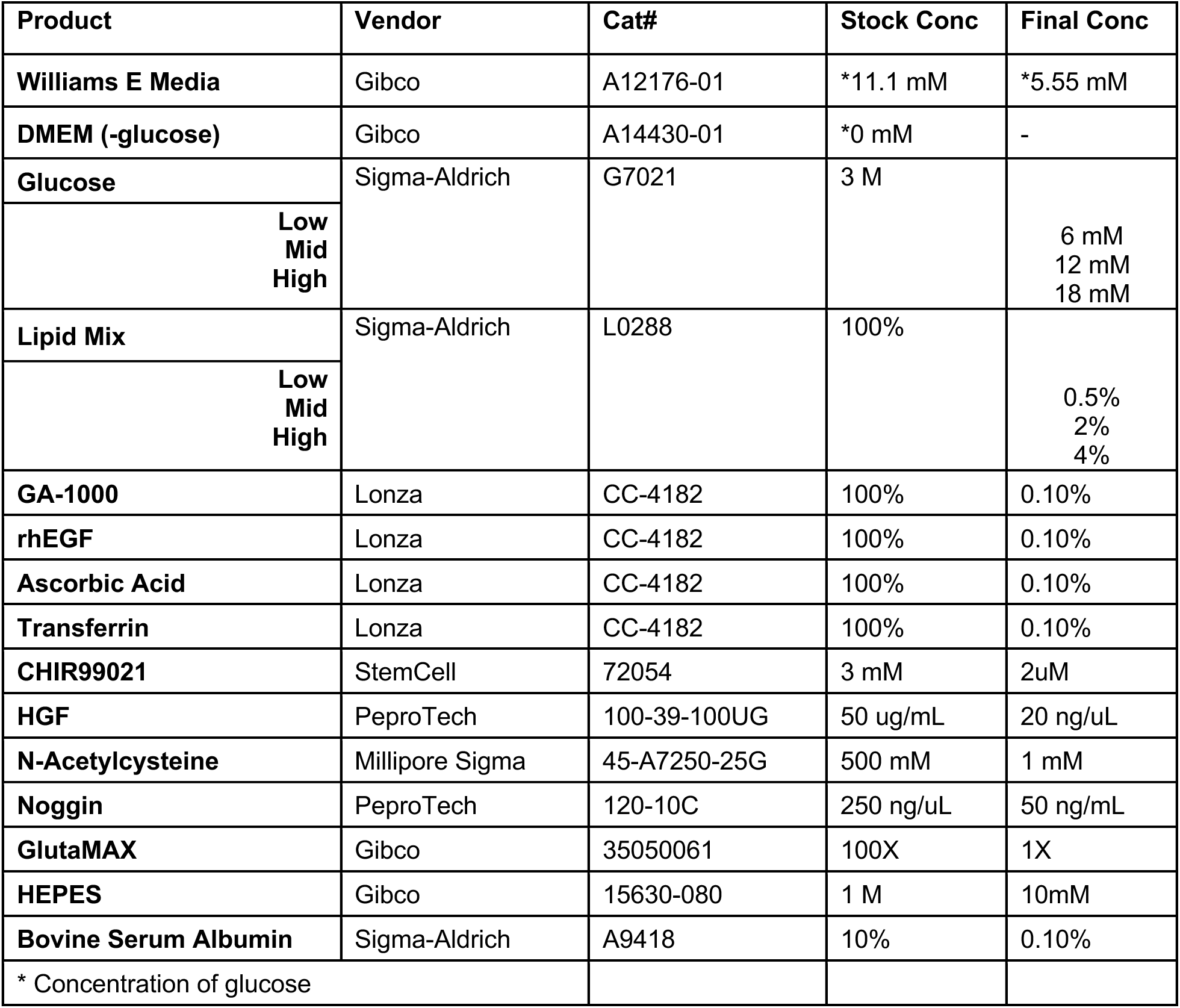
Media formulation.

**Supplementary Table S2:** Lipid Mix 1 formulation.

| **Component** | **Stock Lipid Mixture 1** | **Low (0.5%)** | **Mid (2%)** | **High (4%)** |
| --- | --- | --- | --- | --- |
| **Arachidonic acid** | 2 µg/mL | 0.01 µg/mL | 0.04 µg/mL | 0.08 µg/mL |
| **Each other listed fatty acid** | 10 µg/mL | 0.05 µg/mL | 0.20 µg/mL | 0.40 µg/mL |
| **Cholesterol** | 0.22 mg/mL | 1.1 µg/mL | 4.4 µg/mL | 8.8 µg/mL |
| **Tween-80** | 2.2 mg/mL | 11 µg/mL | 44 µg/mL | 88 µg/mL |
| **Tocopherol acetate** | 70 µg/mL | 0.35 µg/mL | 1.4 µg/mL | 2.8 µg/mL |
| **Pluronic F-68** | 100 mg/mL | 0.5 mg/mL | 2 mg/mL | 4 mg/mL |
| **Ethanol** | 10% v/v | 0.05% v/v | 0.2% v/v | 0.4% v/v |

**Supplementary Figure 1.**
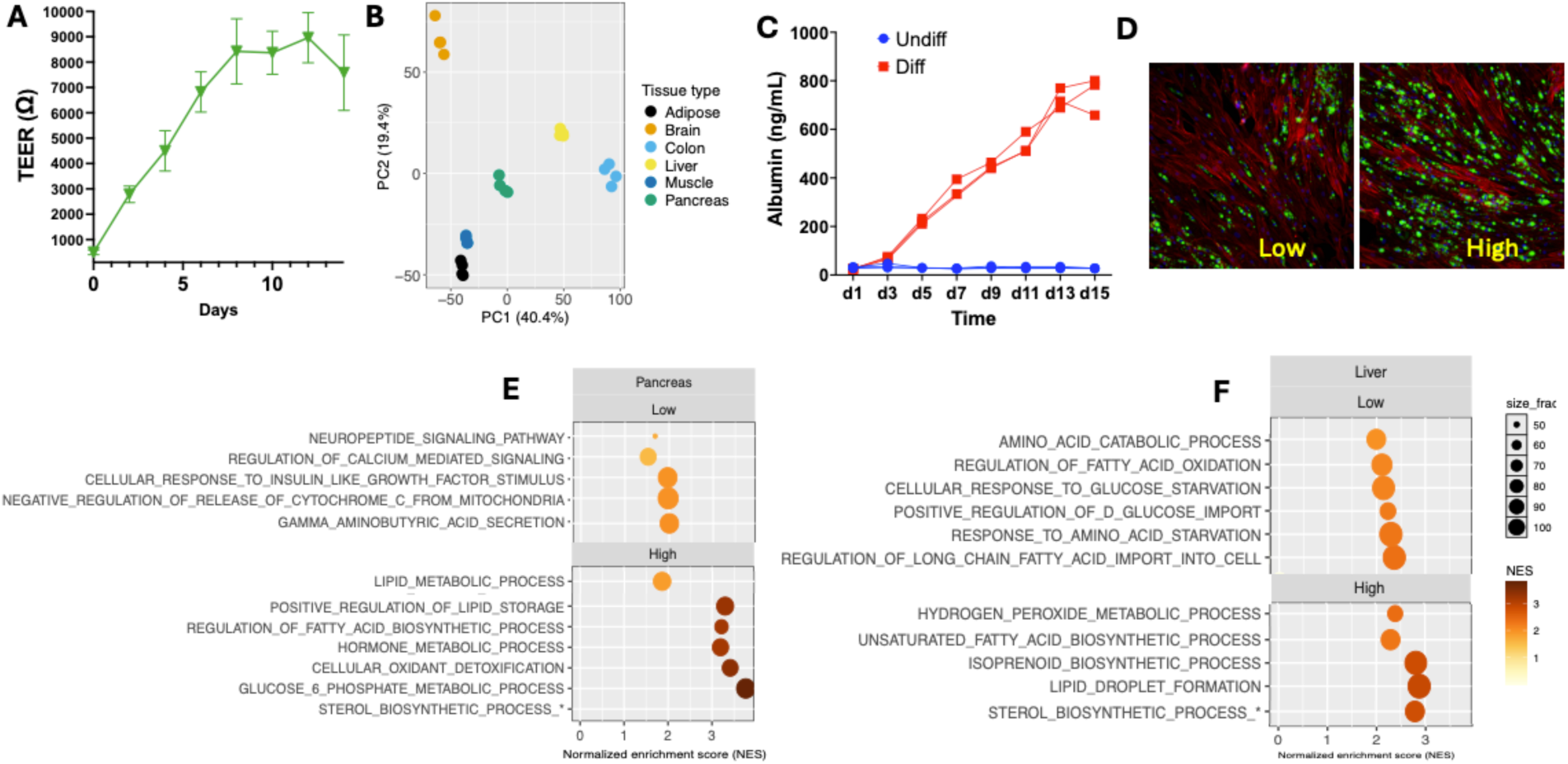
Baseline tissue identity and nutrient responsiveness of isolated tissue modules. **(A)** TEER of gut epithelial monolayers cultured in isolation under basal growth conditions, assessing baseline epithelial barrier formation before multi-tissue interaction experiments. **(B)** Principal component analysis of bulk RNA-seq profiles from six tissue modules cultured in isolation under Mid nutrient conditions. PCA was performed using genes differentially expressed across tissue modules at FDR < 0.05. **(C)** Albumin production by differentiated and undifferentiated liver organoids, supporting acquisition of liver organoid-associated secretory function. **(D)** Representative immunofluorescence images of mature visceral adipocytes cultured under Low or High nutrient conditions. Neutral lipids, green; actin, red; DAPI, blue. Low and High indicate 6 mM glucose/0.5% Lipid Mixture 1 and 18 mM glucose/4% Lipid Mixture 1, respectively. **(E,F)** Selected enriched GO Biological Process pathways in pancreatic islets **(E)** and liver organoids **(F)** cultured in isolation under Low and High nutrient conditions. Dot size represents the number of genes in the data set that overlap with genes within the pathway. Select representative pathways are shown per tissue. RNA-seq data: n = 6 per condition per tissue from two independent experiments.

**Supplementary Figure 2.**
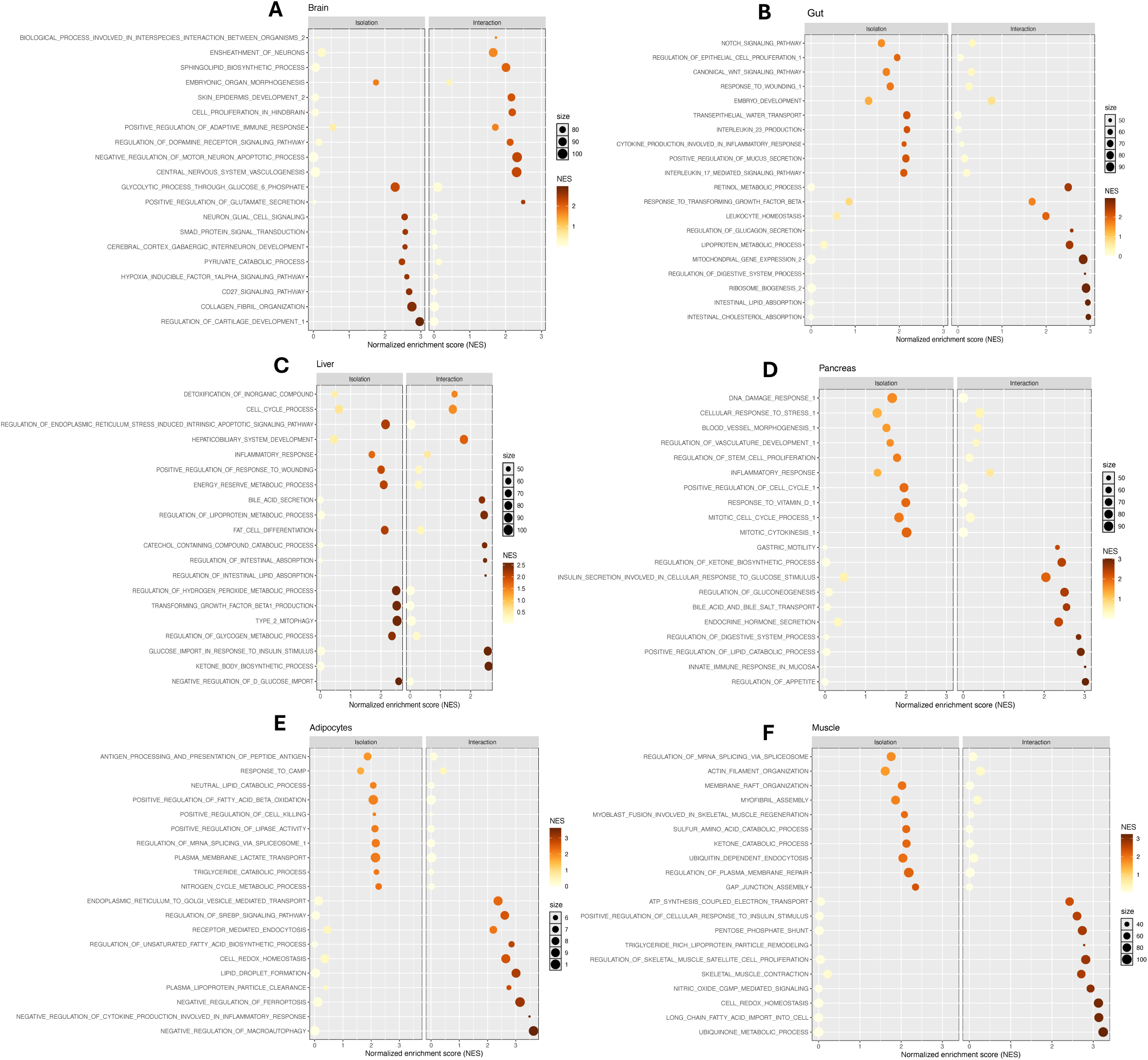
Interaction-associated pathway remodeling across six tissue compartments. GSEA comparing interaction versus isolation under the Mid nutrient condition across **(A)** brain organoids, **(B)** gut, **(C)** liver organoids, **(D)** pancreatic islets, **(E)** adipocytes, and **(F)** skeletal muscle. Mid indicates 12 mM glucose/2% Lipid Mixture 1. Normalized enrichment scores (NES) are shown for selected GO Biological Process pathways; positive NES indicates enrichment in interaction and negative NES indicates enrichment in isolation. Dot size represents the number of genes in the data set that overlap with genes within the pathway. Ten representative pathways are shown per tissue. RNA-seq data: n = 6 per condition per tissue from two independent experiments.

**Supplementary Figure 3.**
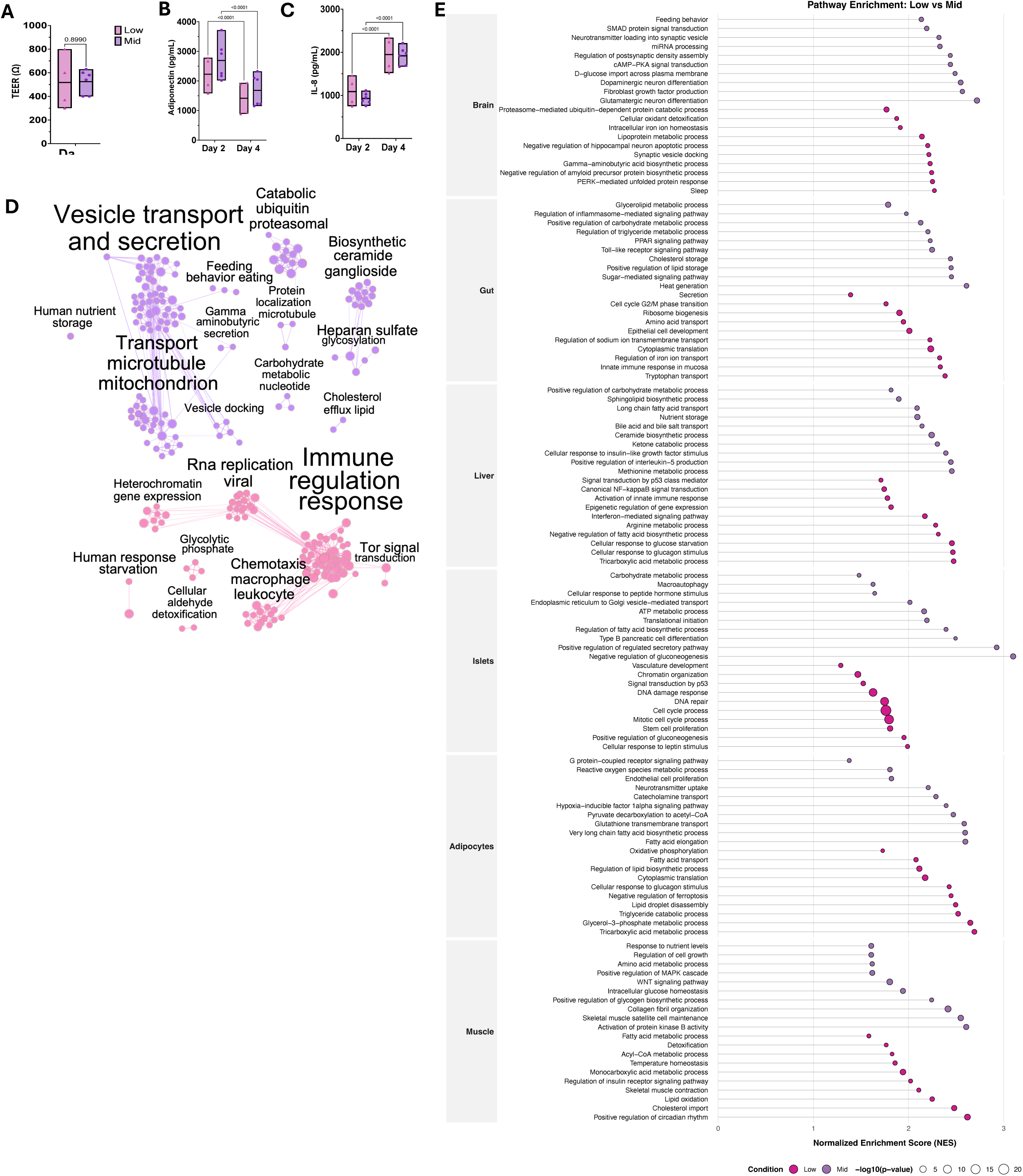
Low-to-Mid nutrient transition remodels epithelial, adipokine, inflammatory, and transcriptional programs in the six-tissue MPS. (A–C) Functional and circulating media readouts from Low and Mid nutrient conditions: (A) TEER of the gut monolayer, (B) adiponectin concentration, and (C) IL-8 concentration. Low and Mid indicate 6 mM glucose/0.5% Lipid Mixture 1 and 12 mM glucose/2% Lipid Mixture 1, respectively. Data are shown as box plots with center line indicating mean and bounds indicating min–max. Statistical significance was determined by two-way repeated-measures ANOVA with nutrient condition and time as factors, followed by Tukey’s multiple-comparisons test. n = 4 for Low and n = 6 for Mid from two independent experiments. (D) Network of significantly enriched GO Biological Process pathways in liver organoids under Low and Mid nutrient conditions, generated from GSEA results using Cytoscape. Node color indicates nutrient condition: Low, pink; Mid, purple. (E) GSEA comparing Low versus Mid nutrient conditions across six tissues. Normalized enrichment scores (NES) are shown for selected GO Biological Process pathways. Color indicates nutrient condition: Mid, purple; Low, pink. Dot size reflects statistical significance as −log₁₀ P value. Ten representative pathways are shown per tissue.

**Supplementary Figure 4.**
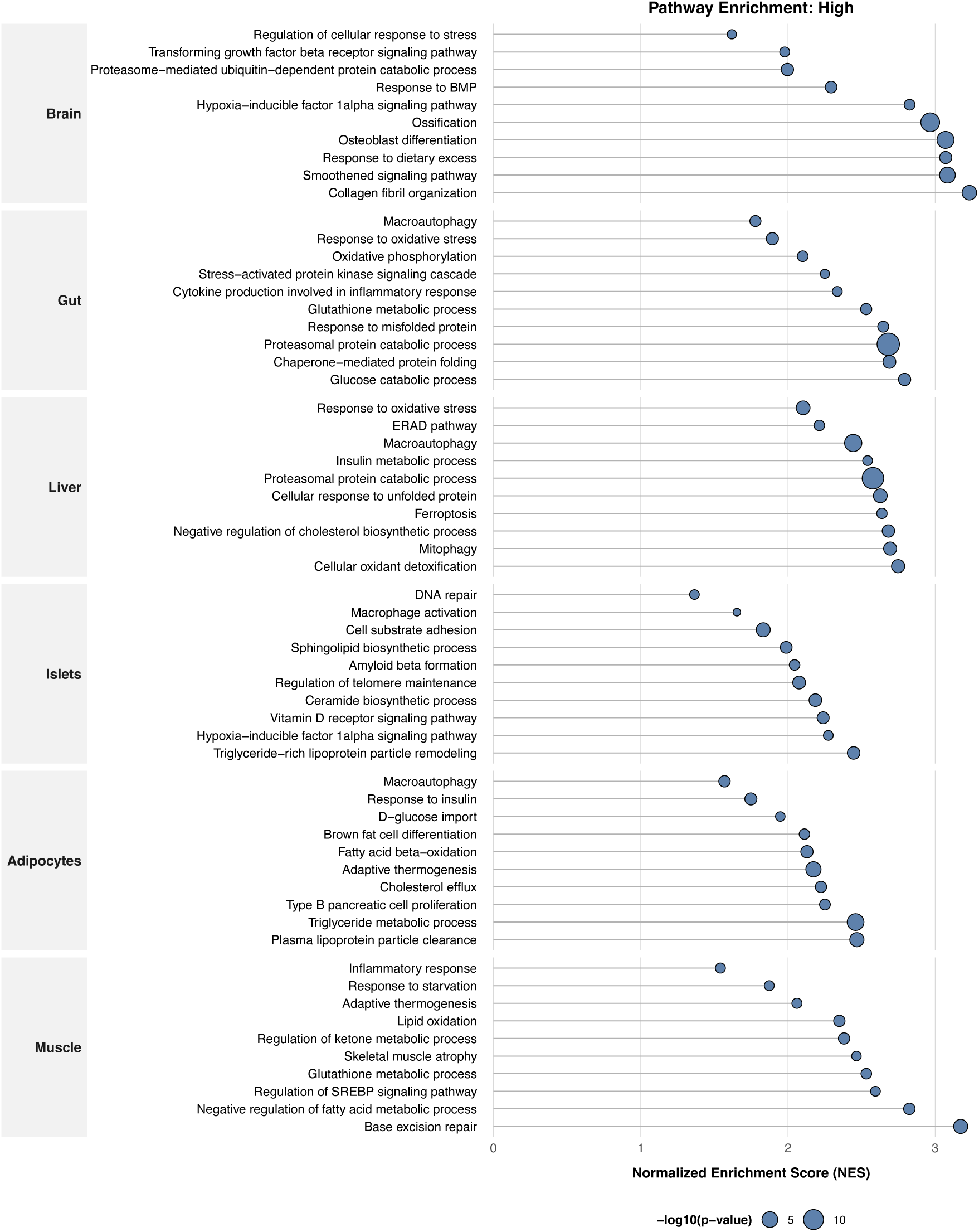
High nutrient exposure remodels stress- and metabolism-associated pathways across the six-tissue MPS. GSEA comparing High versus Mid nutrient conditions across six tissues. High and Mid indicate 18 mM glucose/4% Lipid Mixture 1 and 12 mM glucose/2% Lipid Mixture 1, respectively. Normalized enrichment scores (NES) are shown for selected GO Biological Process pathways. Dot size reflects statistical significance as −log₁₀ P value. Three representative pathways are shown per tissue.

**Supplementary Figure 5.**
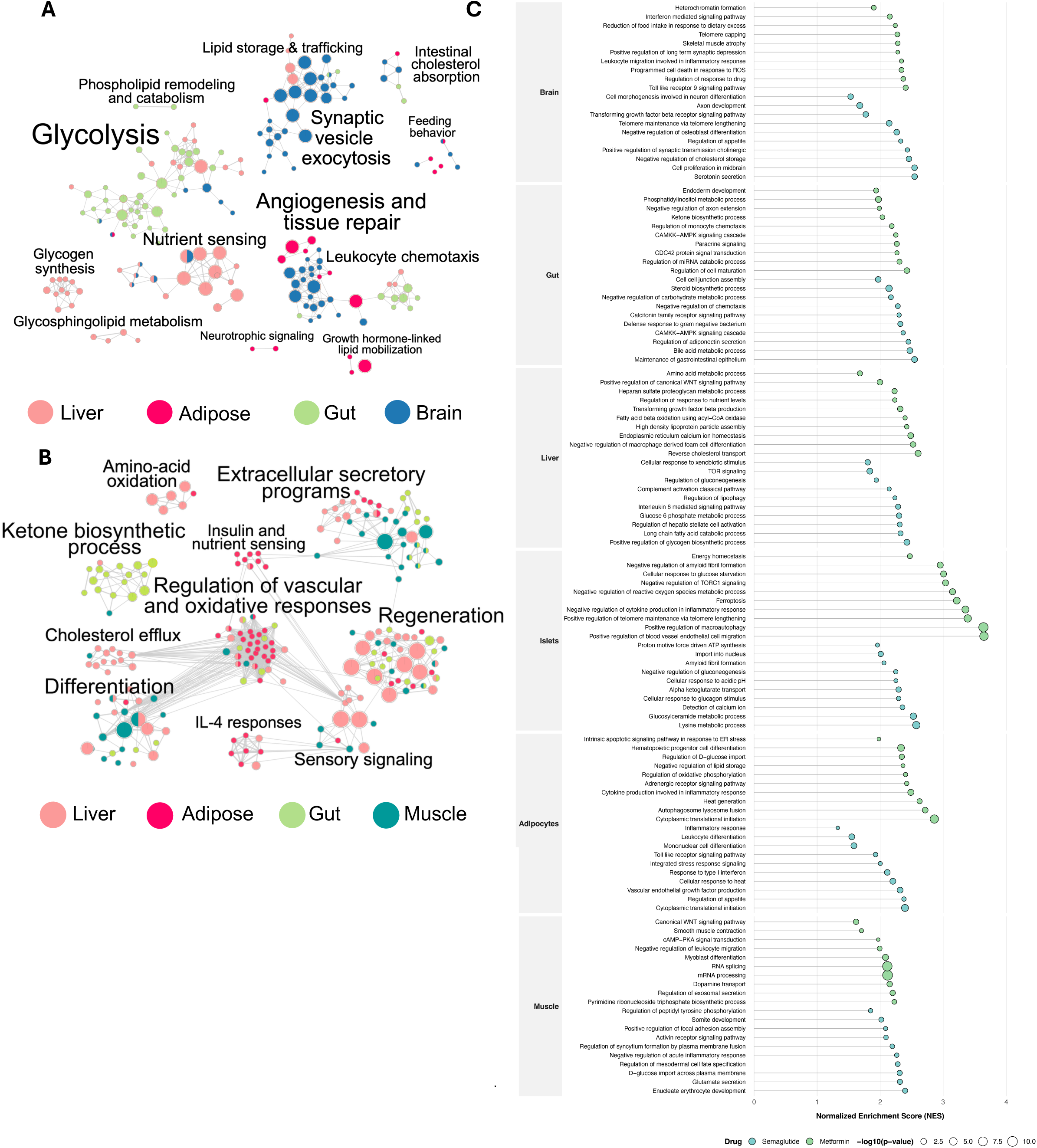
Metformin and semaglutide induce distinct pathway remodeling under High nutrient conditions. **(A)** Semaglutide-induced pathway networks across gut epithelium, liver organoids, adipocytes, and brain organoids under High nutrient conditions, generated from significantly enriched GO Biological Process pathways using Cytoscape. **(B)** Metformin-induced pathway networks across gut epithelium, liver organoids, adipocytes, and skeletal muscle under High nutrient conditions, generated from significantly enriched GO Biological Process pathways using Cytoscape. **(C)** GSEA comparing drug-treated conditions versus untreated High nutrient conditions across six tissues. Normalized enrichment scores (NES) are shown for selected GO Biological Process pathways. Color indicates drug treatment: metformin, green; semaglutide, light blue. Dot size reflects statistical significance as −log₁₀ P value. Three representative pathways are shown per tissue. Significantly enriched pathways in **(A)** and **(B)** were defined as nominal P < 0.05.

